# Zona Incerta subpopulations differentially encode and modulate anxiety

**DOI:** 10.1101/2020.11.11.378364

**Authors:** Zhuoliang Li, Giorgio Rizzi, Kelly R. Tan

## Abstract

Despite recent clinical observations linking the Zona Incerta (ZI) to anxiety, little is known about whether and how the ZI processes anxiety. Here, we subject mice to anxious experiences and observe an increase in ZI cfos-labelled neurons and single-cell calcium activity as well as an efficient effect of ZI infusion of diazepam a classical anxiolytic drug. We further identify that somatostatin-(SOM), calretinin-(CR), and vesicular glutamate transporter-2 (Vglut2)-expressing cells display unique electrophysiological profiles; yet they similarly respond to anxiety-provoking stimuli and to diazepam. Interestingly, optogenetic manipulations reveals that each of these ZI neuronal population triggers specific anxiety-related behavioral phenotypes. Activation of SOM-expressing neurons induced anxiety while in stark contrast, photo-activation of CR-positive cells and photo-inhibition of Vglut2-expressing neurons produce anxiolysis. Furthermore, activation of CR- and Vglut2-positive cells provokes rearing and jumps; respectively. Our findings provide the first experimental evidence that ZI subpopulations encode and modulate different components of anxiety.

## Introduction

Anxiety is a negative emotional state triggered by potential threats and generate autonomic responses, avoidance and stress-like behaviors (*1*, *2*). Several brain regions, including the amygdala (*3*–*5*), the bed nucleus of stria terminalis (*5*, *6*), the medial prefrontal cortex (*7*, *8*), the hippocampus (*4*, *8*, *9*), and the hypothalamus (*9*, *10*) have been reported to play an important role in anxiety. Very recently, the understudied Zona Incerta (ZI) has also been associated with it. Specifically, Parkinson’s disease patients treated with deep brain stimulation (DBS) in the ZI self-reported lower levels of anxiety (*11*, *12*). However, there is little to no experimental evidence supporting these interesting clinical observations.

Located between the thalamus and the hypothalamus, the ZI is an elongated string-like structure that has been implicated in a wide range of functions, varying from neuronal development (*13*), hormonal regulation (*14*, *15*), ingestion (*16*–*18*), sleep (*19*, *20*), sensory processing (*21*, *22*) and pain (*23*, *23*). More recently, a stream of research showcased the implication of the ZI in emotionally related experiences and adaptive defensive behaviors. Exposure to stressful events such as sleep deprivation was shown to potentiate the expression of the immediate early gene cfos in the ZI (*20*). Furthermore, electrical stimulation in the ZI induced pupil dilation (*24*), increased the blood flow to the extremities (*24*) and the heart rate (*25*); all of these are physiological reactions similar to that of the acute stress response (*2*). In addition, ZI loss of function impaired mice in their ability to respond to cues predictive of incoming threat. Specifically, mice with ZI electrolytic lesions displayed reduced avoidance in a conditioned avoidance response test (*26*). Similarly, ZI tetanus toxin infusion to block synaptic transmission decreased the ability to associate a predictive cue with the negative outcome in a fear conditioning task for mice (*27*). In contrast, optogenetic inactivation of ZI Vgat-expressing cells during the presentations of loud tones exacerbated the triggered flight response (*28*). These observations imply that the ZI can sense incoming threats and guide adaptive behaviors to aversive predictive cues. However, how the ZI senses stressful stimuli when mice experience anxiety triggering states is not known.

Most of the current knowledge on the ZI was gained from non-specific manipulations, yet the ZI is a highly heterogenous structure. Indeed, histological investigations revealed diverse cell types possessing the capacity to release chemical signaling molecules such as glutamate, GABA, dopamine or neuropeptides (*29*, *30*). While the GABAergic neurons are abundant in number and thoroughly expressed across the ZI, the existence of somatostatin (SOM), calretinin (CR), calbindin (CB), parvalbumin (PV), and vasopressin (VIP) indicate diversity within the GABAergic cells (*29*–*31*). Such biochemical heterogeneity may explain why the ZI has been implicated in quite a wide range of functions (*30*).

Here we aimed at investigating how the ZI neuronal population responds to stressful and anxious events. We first show that the ZI indeed encodes anxiety. We applied a three-steps strategy with histological labeling against cfos and *in-vivo* calcium recordings as mice experience anxious conditions; as well as the assessment of the efficacy of a pharmacological anxiety-relieving drug directly infused in the ZI. We then focused on two GABAergic subpopulations, the SOM- and CR-expressing neurons, along with the glutamatergic ones. They exhibit distinct electrophysiological properties. Although they are all activated by exposure to anxious conditions and are similarly modulated by diazepam, their bidirectional optogenetic modulation revealed distinct behavioral phenotypes. Specifically, activation of SOM-expressing cells induced anxiogenesis. Conversely, activation of CR-expressing neurons triggered anxiolysis and exploratory rearing. Finally, the activation of Vglut2-expressing cells induced escape-like jumps. Our data provide unprecedented information about how the ZI and some of its neuronal subpopulations encode anxiety related behaviors. Our study hence represents an important first step into understanding the neuronal and circuit basis of the clinical observation for ZI DBS to relieve anxiety in PD patients (*11*, *12*).

## Results

### ZI encodes anxiety

Based on the recently reported clinical observations (*11*, *12*), we first verified and assessed that the ZI encodes anxiety. We performed three experiments among different levels of investigation. First, if indeed the ZI encodes anxiety, an anxious stimulus such as being forced to stay on an elevated platform is expected to increase the expression of the immediate early gene cfos in the ZI. Mice were hence placed on an elevated circular platform (100cm above ground, 3 cm in diameter, Figure 1A) for 30 minutes and brought back to their home cage for an additional 30 min before intracardiac perfusion. Control mice stayed in their home cage for 60 min (Figure 1B). Post-hoc immunostaining against cfos (Figure 1C-D) followed by quantification revealed a significantly higher number of cfos labelled neurons in the ZI of mice which experienced the elevated platform as compared to control mice (Figure 1E, elevated rod 3.38 ± 0.20 vs home cage 1.00 ± 0.14 cells, 951 and 243 cells respectively in 12 slices from 3 mice). These data confirm that an anxiety or stress triggering event increased the activity of ZI cells.

**Figure 1.**
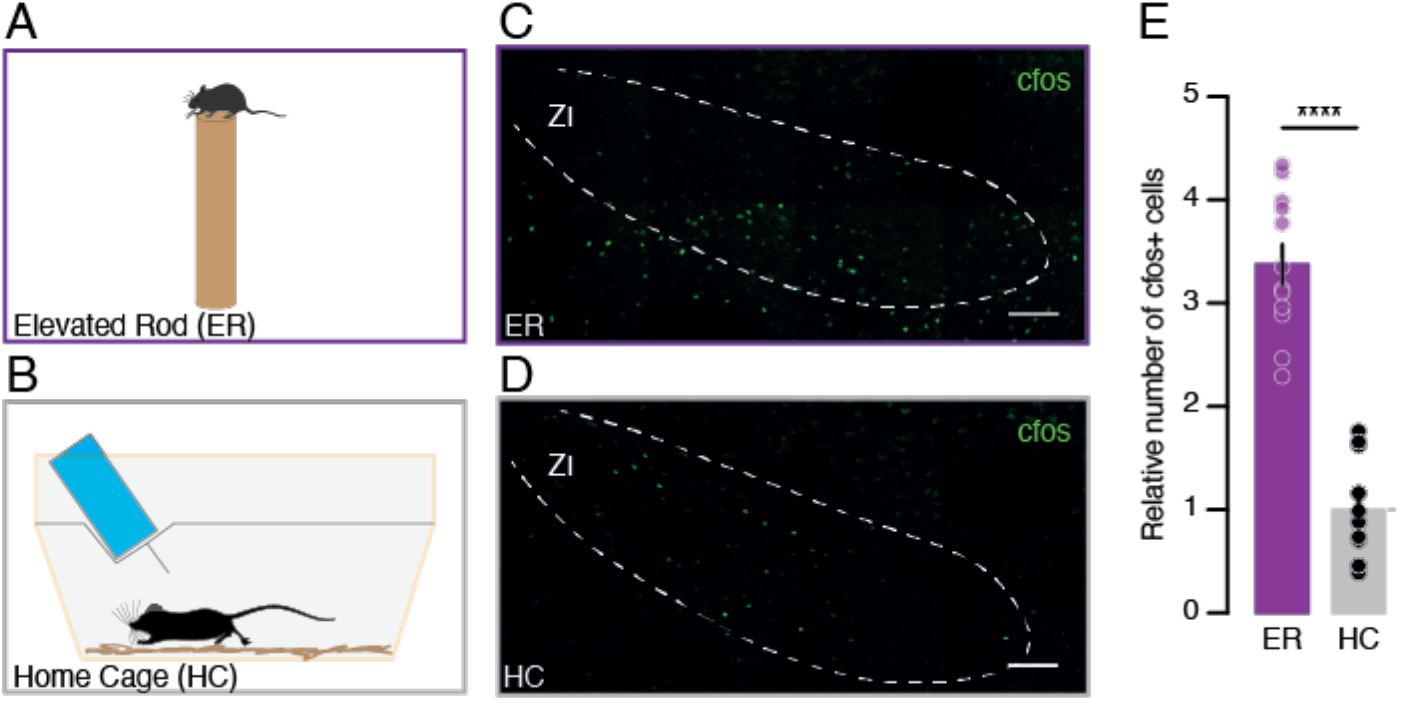
Anxious experience enhances ZI neuronal activation. A) Experimental setup where mice were placed on a narrow-elevated rod. B) Schematic of the control home cage condition. C) Example confocal image of the immunohistochemical (IHC) staining against cfos (green) in the ZI-containing coronal brain slices from mice that were exposed to the anxious elevated rod experience (100μm scale bar). D) Example confocal image of the IHC labeling of cfos in the ZI-containing coronal brain slices of mice that were left in the control home cage condition (100μm scale bar). E) Bar graphs reporting the relative number of cfos labeled cells (normalized to the control mice) in the ZI for both experimental groups. Data are presented with the mean ± SEM.

We next further investigated this with a higher temporal precision through *in vivo* calcium imaging. An AAV construct carrying the calcium indicator GCaMP6m gene was expressed in ZI neurons and a GRIN lens implanted slightly above (Figure 2A-B). We recorded the calcium dynamics of ZI neurons while mice explored an elevated platform inspired from the elevated plus maze. Such experimental apparatus allowed us to acquire and analyze calcium transients in ZI neurons while animals freely explored between the safe center and the anxiety-evoking edges of the open arms (Figure 2C). We observed that a significant proportion of cells exhibited higher levels of activity when mice explored the safe center while other cells showed elevated calcium activity in the open edges (Figure 2D-E, Video S1). To confirm and strengthen such observation we further analyzed and evaluated the correlation between neuronal activity with the absolute distance of the recorded animal away from the safe center. 23% of the imaged ZI cells displayed a positive correlation and were more active when mice explored the anxious edges as compared to the safer center (Figure 2F-H, +ve Corr 0.96 ± 0.07 vs not Corr 0.06 ± 0.04 z-scores, 128 cells from 5 mice). Conversely, 9% of the imaged cells exhibited a negative correlation and were less active when the animals were in the edges compared to the center (Figure 2F-H, −ve Corr −0.75 ± 0.06 vs not Corr 0.06 ± 0.04 z-scores). We also analyzed the calcium dynamics as mice transitioned into a safer or more anxious location (Figure 2I-L). To do so, we extracted from the cells exhibiting a positive, negative and no correlation (from Figure 2F-H), the calcium activity immediately before and after mice transitioned into and out of the edges (edge entry and exit; respectively, Figure 2I-J) as well as the center (center exit and entry, Figure 2 K-L). The positively correlated neurons increased their activity when entering the edge or leaving the center (Figure 2I, edge entry, center/transition 0.00 ± 0.07 vs edge 0.32 ± 0.06 z-scores; Figure 2K, center exit, center 0.00 ± 0.03 vs edge/transition 0.15 ± 0.04 z-scores) and significantly decreased their activity when leaving the edge or entering the center (Figure 2J, edge exit, edge 0.00 ± 0.10 vs center/transition −0.36 ± 0.03 z-scores; Figure 2L, center entry, edge/transition 0.00 ± 0.03 vs center −0.20 ± 0.05 z-scores). On the other hand, the negatively correlated cells showed the opposite trend; reaching significant decreases in activity when the animals exited the edges (Figure 2I, edge entry, center/transition 0.00 ± 0.06 vs edge - 0.259 ± 0.03 z-scores; Figure 2J, edge exit, edge 0.00 ± 0.03 vs center/transition 0.18 ± 0.07 z-scores; Figure 2K, center exit, center 0.00 ± 0.03 vs edge/transition −0.07 ± 0.04 z-scores; Figure 2L, center entry, edge/transition 0.00 ± 0.02 vs center −0.06 ± 0.02 z-scores). Corresponding increases and decreases in activity when mice entered more anxiety provoking and safer areas; respectively, provide further support that this subset of ZI neurons encode anxiety related information. To exclude a possible motor component, we correlated the calcium activity of individual ZI cells with the movement speed of the recorded mice. Of the cells that displayed positive and negative correlation with the mouse position on the platform; a majority of 69 and 91% were not correlated with the movement speed; respectively (Figure S1A-B, 20 of 29 and 10 of 11 cells). On average, the speed mice traveled in the center and the edges were similar (Figure S1C, center 0.038 ± 0.003 vs edges 0.024 ± 0.007 m/s) and thus further suggest that movement speed does not explain the differences in neuronal activity while mice were in these two areas. All together, these data reveal that a significant subset of ZI neurons is sensitive to anxiety-related cues irrespective of locomotion.

**Figure 2.**
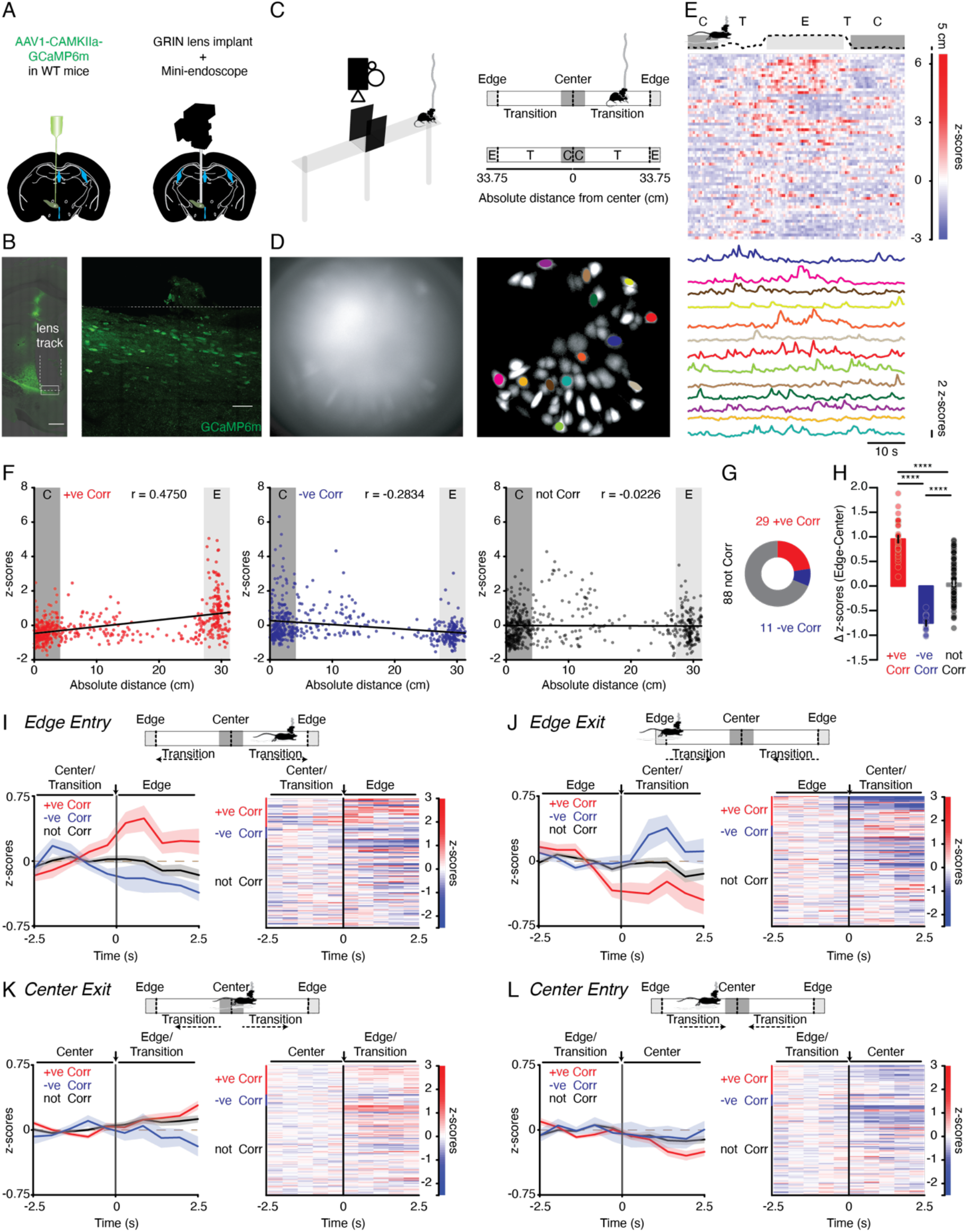
ZI neurons are sensitive to anxiety-related cues. A) Surgical setup to express and visualize GCaMP6m in the ZI of wildtype mice. B) Example confocal image of a partial coronal section containing the GRIN lens tract and the GCaMP6m expressing (green) cells in the ZI (500 and 50μm scale bar, respectively). C) Schematic of the elevated platform design, viewed from the side (left) and from above (right) with the compartments (edge, transition, and center) annotated along with the distance from the center to edges. D) Example frame in the raw ZI calcium imaging recording video (left) and maximum intensity projection visualization of identified cells following data pre-processing (right) with example cells color coded. E) Track plot of the center of a single mouse (top) as it traveled along the elevated platform with the different compartments highlighted. Time-locked heatmap of the calcium activity (represented in z-scores) of all recorded cells (middle) and traces from example cells (bottom; corresponding to ones labeled in Figure 2E) from the position-tracked mouse. F) Scatterplots of example recorded cells with calcium activity positively (red), negatively (blue), and not (grey) correlated to the distance of the recorded mouse from the center on the elevated platform. G) Pie chart depicting the proportion of cells whose calcium activity positively, negatively, and not correlated to the center distance. H) Bar graphs reporting the differences in calcium activity while mice were in the edges and in the center between the three types of center distance correlated cells. I) Transient changes in activity of the three types of center distance correlated cells as mice entered the edge. Schematic of the positional change of the recorded mice from the center/transition area to the edge (top). Line plot (left) depicting the average activity of center distance correlated cells 2.5s before and after the positional change (align at 0s, denoted by the arrow). Heatmap (right) of the activity of all cells (grouped and sorted based on their correlation coefficient) before and after positional change. J) Transient changes in activity of the three types of center distance correlated cells as mice exited the edge. K) Transient changes in activity of the three types of center distance correlated cells as mice exited the center. L) Transient changes in activity of the three types of center distance correlated cells as mice entered the center. Data are presented with the mean ± SEM.

To assess the ability of the ZI to influence anxiety, we took a pharmacological approach where the anxiolytic properties of diazepam applied directly into the ZI was tested in the elevated O-maze (EOM, Figure 3). Infusion cannulas were bilaterally implanted above the ZI (Figure 3A-B). Diazepam (1.5mg/kg) or vehicle infusions were made 30 min prior to placing mice in the EOM (Figure 3C). The order of the treatments was counterbalanced and reversed 2 weeks later. When mice received diazepam, they spent significantly more time in the open arms as compared to when they were treated with the vehicle (Figure 3D-E, Video S2, vehicle 130.8 ± 27.85 vs diazepam 219.9 ± 30.03s, 13 mice). They also made more entries into the open arms (Figure 3F, vehicle 24.38 ± 4.90 vs diazepam 37.92 ± 4.77 entries). Such differences are not due to diazepam-induced motor effects since the distance traveled and the time mice spent immobile in the open field test (OFT) were similar with both treatment conditions (Figure 3G-I, distance traveled, vehicle 43.76 ± 3.82 vs diazepam 46.33 ± 3.22; time immobile vehicle 50.05 ± 13.17 vs diazepam 64.39 ± 17.09s). These findings suggest that local diazepam treatment in the ZI reduces anxiety without impacting locomotion, further consolidating our biochemical and *in vivo* calcium imaging data. To briefly summarize here, our multi-level experimental observations converge onto the ZI as a brain structure processing anxiety-related stimulus.

**Figure 3.**
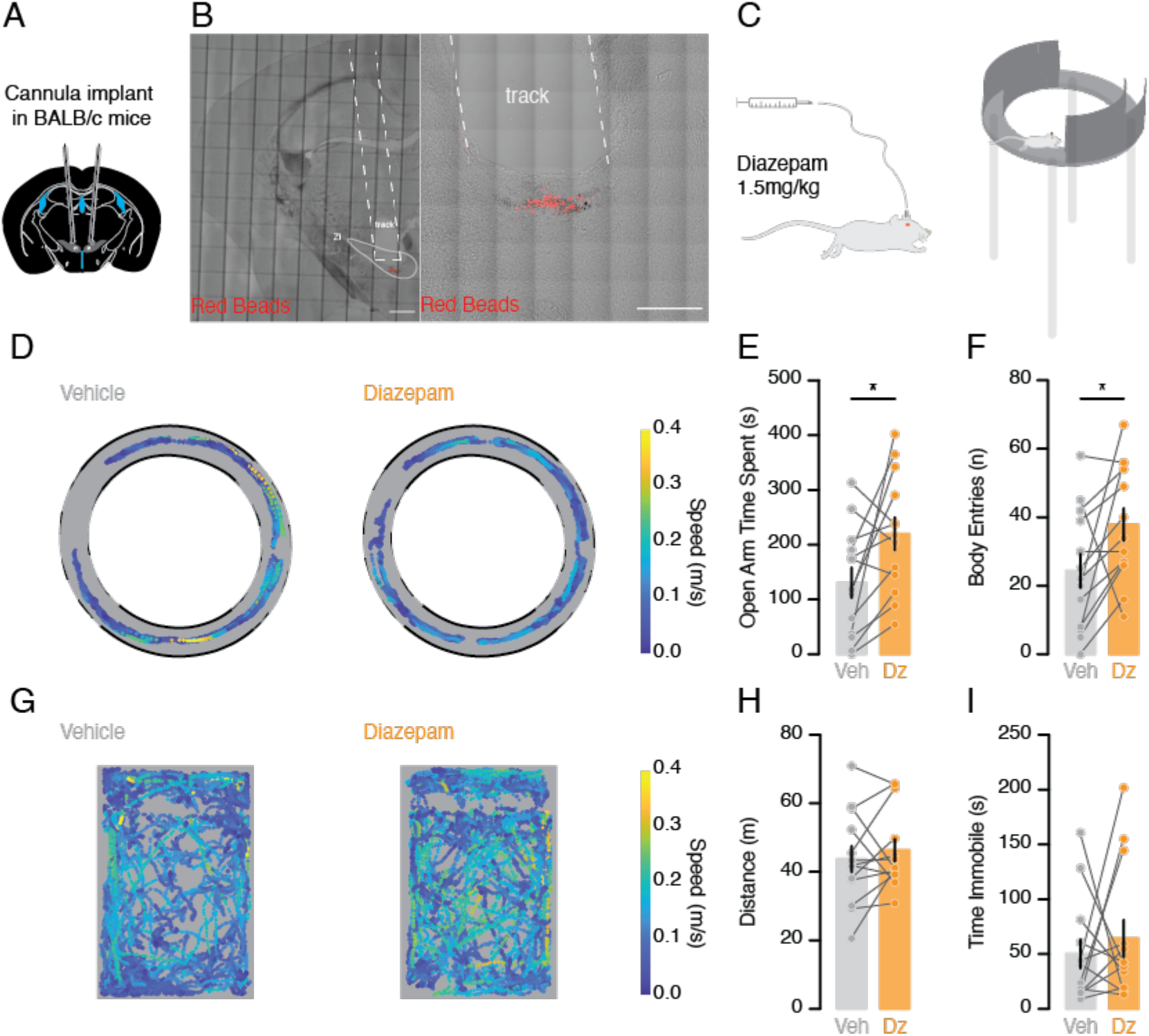
Local ZI diazepam infusion reduces anxiety without affecting movement. A) Surgical setup to place cannulas above the ZI. B) Example confocal image of a partial coronal section containing the cannula tract with the red retrobeads in the ZI to show the extent of spread (500 and 250μm scale bar, respectively). C) Schematic of the setup to directly infuse diazepam (1.5mg/kg) into the ZI or vehicle 30 minutes prior to testing the mouse on the EOM. D) Track plot of a single example mouse on the EOM (dashed lines denote the open arms while the filled lines indicate the closed arms) after vehicle (left) and diazepam (right) infusion with the speed color-coded. E) Bar graph of time the mice spent in the open arms after vehicle and diazepam infusions. F) Bar graph depicting the number of times mice entered the open arm after vehicle and diazepam infusions. G) Track plot of an example mouse in the OFT after vehicle (left) and diazepam (right) infusion with the speed color-coded. H) Bar graph depicting the distance mice traveled after vehicle or diazepam infusions. I) Bar graph showing the time mice spent immobile after vehicle or diazepam infusions. Data are presented with the mean ± SEM.

### ZI SOM-, CR- and Vglut2-expressing cells exhibit unique electrophysiological profiles and are all sensitive to anxious stimuli and diazepam

So far, we have shown with complementary approaches that the ZI encodes anxiety. However, it is important to note that we observed subsets of ZI neurons displaying anxiety-induced cfos expression and anxiety-induced changes in calcium transients. Previous literature reported the cellular diversity within the ZI (*29*–*31*). We therefore wondered about the specific role an individual ZI subpopulation plays in the encoding of anxiety. To address this question, we first examined the ZI biochemical diversity with FISH, focusing on the GABAergic and glutamatergic markers with Vgat and Vglut2; respectively. We report that Vgat-expressing cells are more abundant than Vglut2-expressing ones (Figure 4A-B; Vgat-only 78.8% and Vglut2-only 19.0%, 3390 cells from 3 mice). Probing against specific inhibitory markers namely somatostatin (SOM), calretinin (CR), and parvalbumin (PV) revealed that the SOM- and CR-expressing subpopulations are larger than the PV-expressing one (Figure 4C-D; SOM-only 36.6%, CR-only 53.2%, PV-only 2.0%, 1684 cells from 3 mice). Expectedly, SOM- and CR-expressing cells co-express Vgat (Figure S2A-D). In the rest of our study, we focused on the SOM- and CR-as well as the Vglut2-expressing cells.

**Figure 4.**
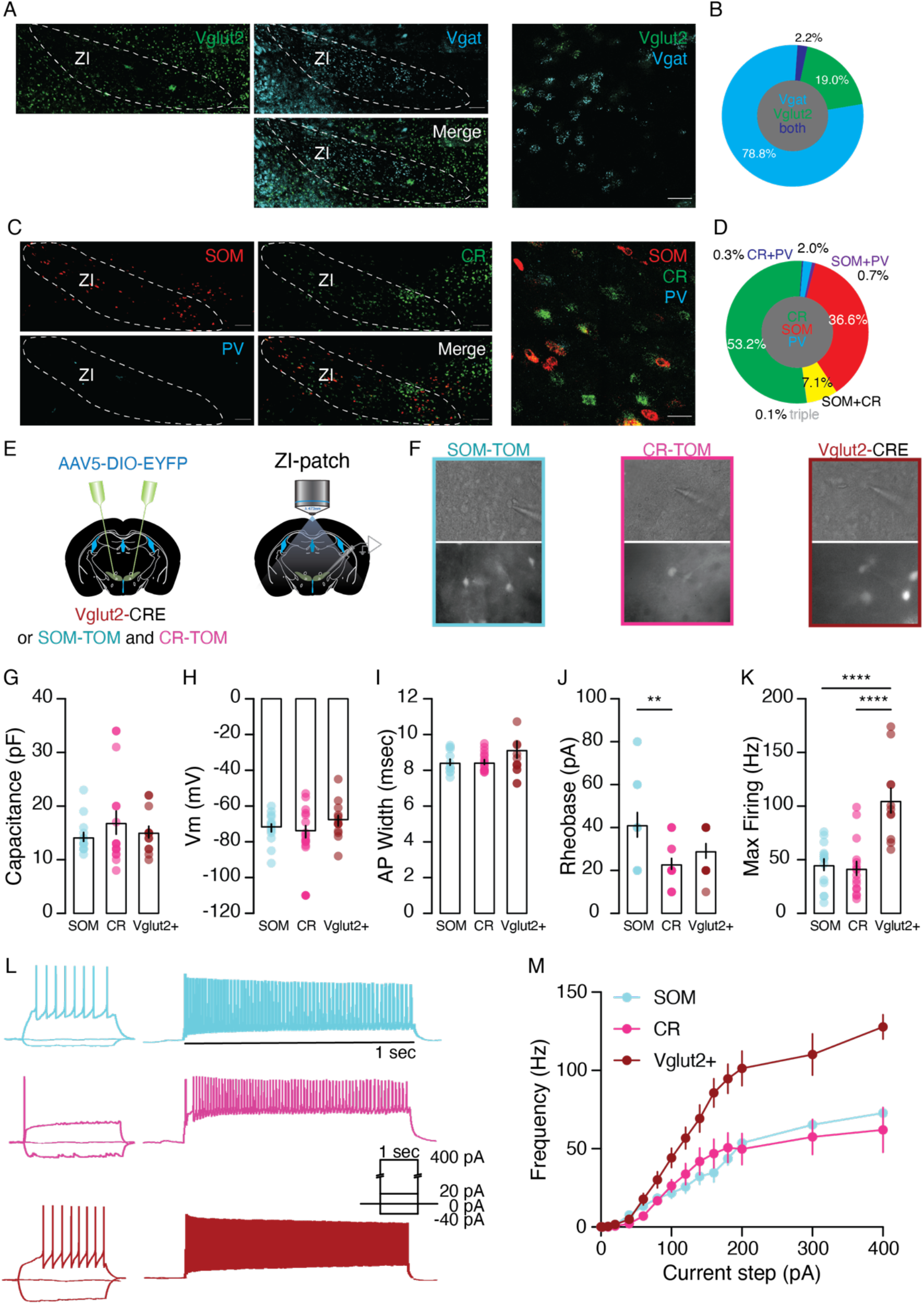
SOM-, CR-, and Vglut2-expressing ZI neurons display unique electrophysiological properties. A) Example confocal images of ZI-containing coronal brain slices with the FISH labeling of the mRNAs for Vglut2 (green) and Vgat (cyan) as well as the merged image (100μm and 25μm scale bar respectively). B) Pie chart quantification of the proportion of cells in ZI expressing Vglut2, Vgat, or both C) Example confocal images of the ZI-containing coronal brain sections with the FISH labeling of the mRNAs for SOM (red), CR (green), PV (cyan), and the merged (100μm and 25μm scale bar respectively). D) Pie chart representation of the percentage of expressing the mRNAs for SOM, CR, PV, or any combination of the three. E) Schematic of the experimental procedure. F) Example brightfield (top) and fluorescent (bottom) images of patched cells: SOM (cyan), CR (magenta), and Vglut2 (maroon). G) Bar graphs reporting the capacitance from SOM-, CR-, and Vglut2-expressing ZI cells. H) Bar graphs depicting the resting membrane potential (Vm) from SOM-, CR-, and Vglut2-expressing ZI cells. I) Bar graphs of the action potential width from SOM-, CR-, and Vglut2-expressing ZI cells. J) Bar graphs presenting the rheobase from SOM-, CR-, and Vglut2-expressing ZI cells. K) Bar graphs summarizing the maximum firing frequency from SOM-, CR-, and Vglut2-expressing ZI cells. L) Example current clamp recordings from SOM-, CR-, and Vglut2-expressing ZI cells upon various current step injections. M) Summary input/output curve for SOM-, CR-, and Vglut2- positive cells. Data are presented with the mean ± SEM.

Next, we functionally characterized these ZI subpopulations performing *in-vitro* whole cell patch clamp recordings (Figure 4 E-M). To unequivocally study SOM-, CR-, and Vglut2-expressing cells, reporter lines (SOM-TOM and CR-TOM) or CRE-dependent EYFP (AAV5-DIO-EYFP) injected Vglut2-CRE-mouse line were used (Figure 4E-F). Despite exhibiting similar capacitance (Figure 4G, SOM 14.27 ± 0.80, CR 16.94 ± 2.16, Vglut2 15.15 ± 1.09 pF, 13-16 cells, 4-5 mice), resting membrane potential (Figure 4H, SOM −72.20 ± 2.12, CR −74.31 ± 3.37, Vglut2 −68.15 ± 2.88 mV), and action potential width (Figure 4I, SOM 8.46 ± 0.16, CR 8.46 ± 0.13, Vglut2 9.16 ± 0.47ms), SOM-expressing neurons are less excitable and display a higher rheobase (Figure 4J, SOM 41.33 ± 5.68, CR 23.13 ± 2.54, Vglut2 29.17 ± 3.362 pA) while Vglut2-expressing cells sustain higher maximum firing frequencies (Figure 4K-M, SOM 45.43 ± 5.34, CR 42.00 ± 6.38, Vglut2 105.20 ± 11.42 Hz).

We next assessed if SOM-, CR-, and Vglut2-expressing ZI neurons are activated by anxious conditions. The elevated rod experiment followed by cfos labeling was performed (Figure S3). Our quantification reports that co-expression of cfos with either SOM-, CR- or Vglut2-is significantly increased in the experimental mice as compared to the control ones (Figure S3B-P, SOM: elevated rod 3.50 ± 0.35 vs home cage 1.00 ± 0.18 cells, 3262 and 2458 cells total respectively in 12 slices from 3 mice; CR: elevated rod 7.40 ± 1.39 vs home cage 1.00 ± 0.25 cells, 2074 and 1624 cells total, respectively; Vglut2: elevated rod 3.28 ± 0.30 vs home cage 1.00 ± 0.17 cells, 2538 and 1551 cells total, respectively). These data suggest that SOM-, CR-, and Vglut2-expressing cells are all activated by anxiety.

Finally, we assessed the sensitivity to diazepam in these three neuronal cell types. *In vitro* recordings of pharmacologically isolated inhibitory post-synaptic currents (IPSCs) onto SOM-, CR- and Vglut2-expressing cells were performed. The neurons were identified through the expression of TdTomato/EGFP (Figure 5A). Diazepam (1μM) potentiated the electrically evoked-IPSCs (Figure 5B, SOM: baseline −523.9 ± 96.29 vs diazepam −907.7 ± 175.3 pA; CR: baseline −232.9 ± 63.65 vs diazepam −500.3 ± 85.55 pA; Vglut2: baseline −342.0 ± 71.26 vs −669.0 ± 106.5 pA; 10-11 cells from 4-5 mice) without affecting the pair-pulse ratio in all three cell types (Figure 5B, SOM: baseline 1.11 ± 0.09 vs diazepam 0.91 ± 0.07; CR: baseline 0.95 ± 0.11 vs diazepam 0.85 ± 0.08; Vglut2: baseline 1.06 ± 0.07 vs diazepam 0.98 ± 0.06). The amplitude (Figure 5C, SOM: baseline −41.78 ± 5.07 vs diazepam −48.65 ± 6.27 pA; CR: baseline −44.54 ± 3.05 vs diazepam −50.08 ± 3.12 pA; Vglut2: baseline 40.79 ± 3.22 vs diazepam −47.54 ± 3.98 pA; 10-12 cells from 4-5 mice) and frequency (Figure 5C, SOM: baseline 0.05 ± 0.02 vs diazepam 0.09 ± 0.03 Hz; CR: baseline 0.04 ± 0.01 vs diazepam 0.06 ± 0.01 Hz; Vglut2: baseline 0.08 ± 0.02 vs diazepam 0.09 ± 0.02 Hz) of the spontaneous IPSCs were both increased. The IPSCs were blocked by picrotoxin (100μM). These *in vitro* electrophysiological data reveal that all three neuronal subpopulations are sensitive to diazepam.

**Figure 5.**
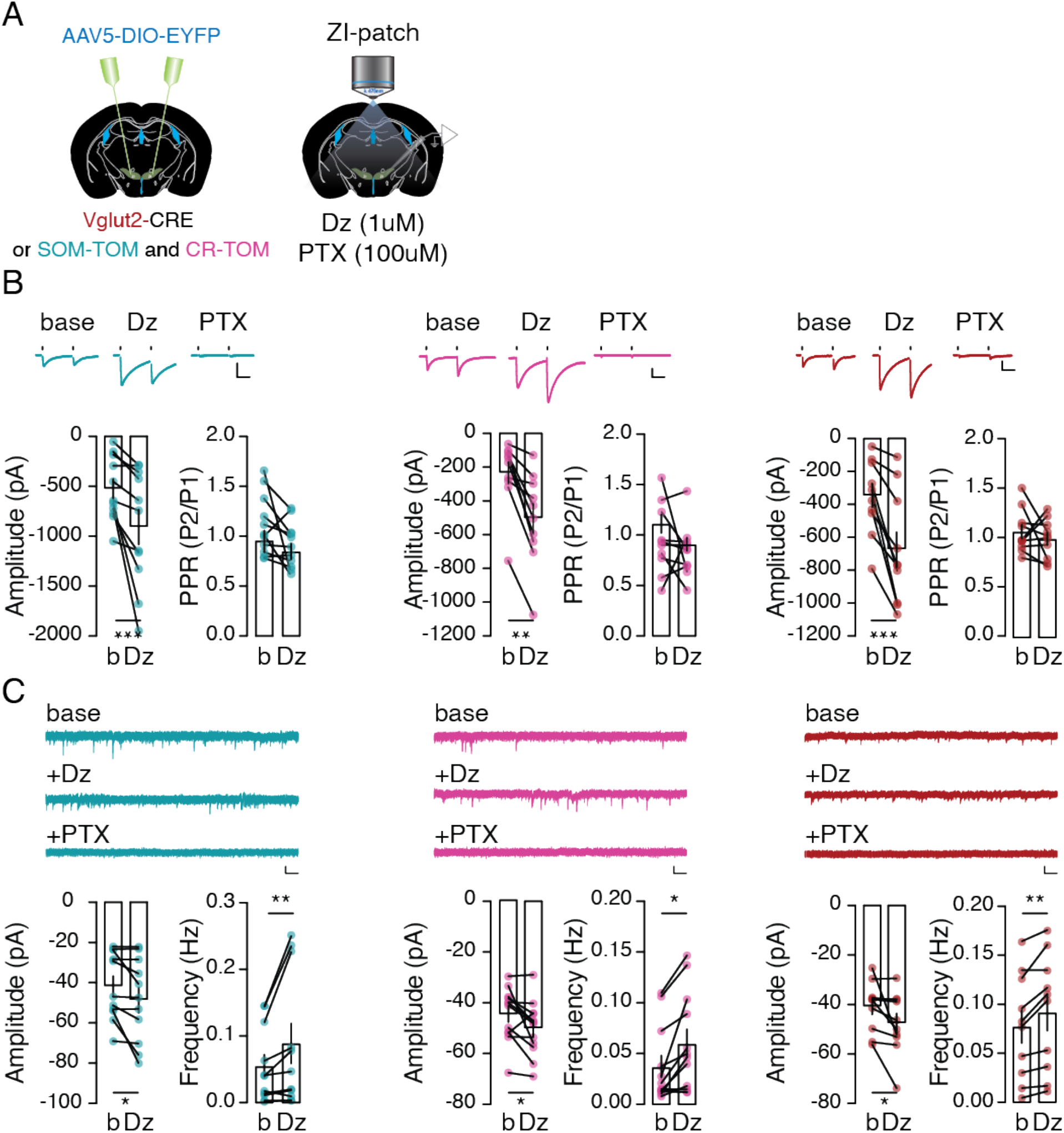
Diazepam potentiates inhibitory drive onto SOM-, CR-, and Vglut2-expressing ZI neurons. A) Schematic of the experimental setup. B) Example IPSC from representative SOM- (cyan), CR- (magenta) and Vglut2- (maroon) expressing ZI cells during baseline (base) and following subsequent bath application of diazepam (Dz) and picrotoxin (PTX). The scale bars represent 100pA (vertical) and 25ms (horizontal). Bar graphs report the amplitude of the IPSCs and the paired-pulse ratio (PPR) both conditions. C) Example sIPSCs recorded from representative SOM-, CR-, and Vglut2-expressing ZI cells during baseline (base) and during bath application of diazepam (Dz) and picrotoxin (PTX). The scale bars represent 30pA (vertical) and 500ms (horizontal). Bar graphs of the amplitude and the frequency of sIPSCs in the baseline and following diazepam application. Data are presented with the mean ± SEM.

Altogether, these findings suggest that SOM-, CR-, and Vglut2-expressing cells of the ZI are distinct neuronal sub-populations with specific functional intrinsic properties. However, they are all similarly activated by anxious stimuli and are sensitive to diazepam.

### SOM-, CR-, and Vglut2-expressing ZI neurons trigger distinct anxiety-related traits

Despite being activated by elevation and allosterically modulated by diazepam; SOM-, CR-as well as Vglut2-expressing cells might still engage different components of anxiety-related behaviors. To verify this hypothesis, we bidirectionally manipulated the activity of these three subpopulations while mice performed in the EOM task (Figure 6). To do so, we injected AAV vectors containing either CRE-dependent ChR2 (AAV5-DIO-ChR2-EYFP) or Arch (AAV5-DIO-Arch-EYFP) in the ZI of SOM-, CR-, and Vglut2-CRE mice and implanted optic fibers above the infection site (Figure 6A-B). Our experimental preparations were validated through *in-vitro* patch clamp recordings (Figure S4A-B). A short blue light illumination (400 msec, 473 nm) evoked desensitizing photocurrents in ChR2-positive SOM-, CR-, and Vglut2-positive ZI neurons (Figure S4C). Furthermore, either a 1 second continuous light pulse or a 10 second (40Hz) stimulation elicited firing activity that was reversed as soon as the stimulation was turned OFF (Figure S4D). In contrast, green light pulses (400 msec, 532nm) triggered hyperpolarizing photocurrents in Arch expressing SOM-, CR-, and Vglut2-positive cells, preventing evoked or spontaneous firing (Figure S4E-F).

**Figure 6.**
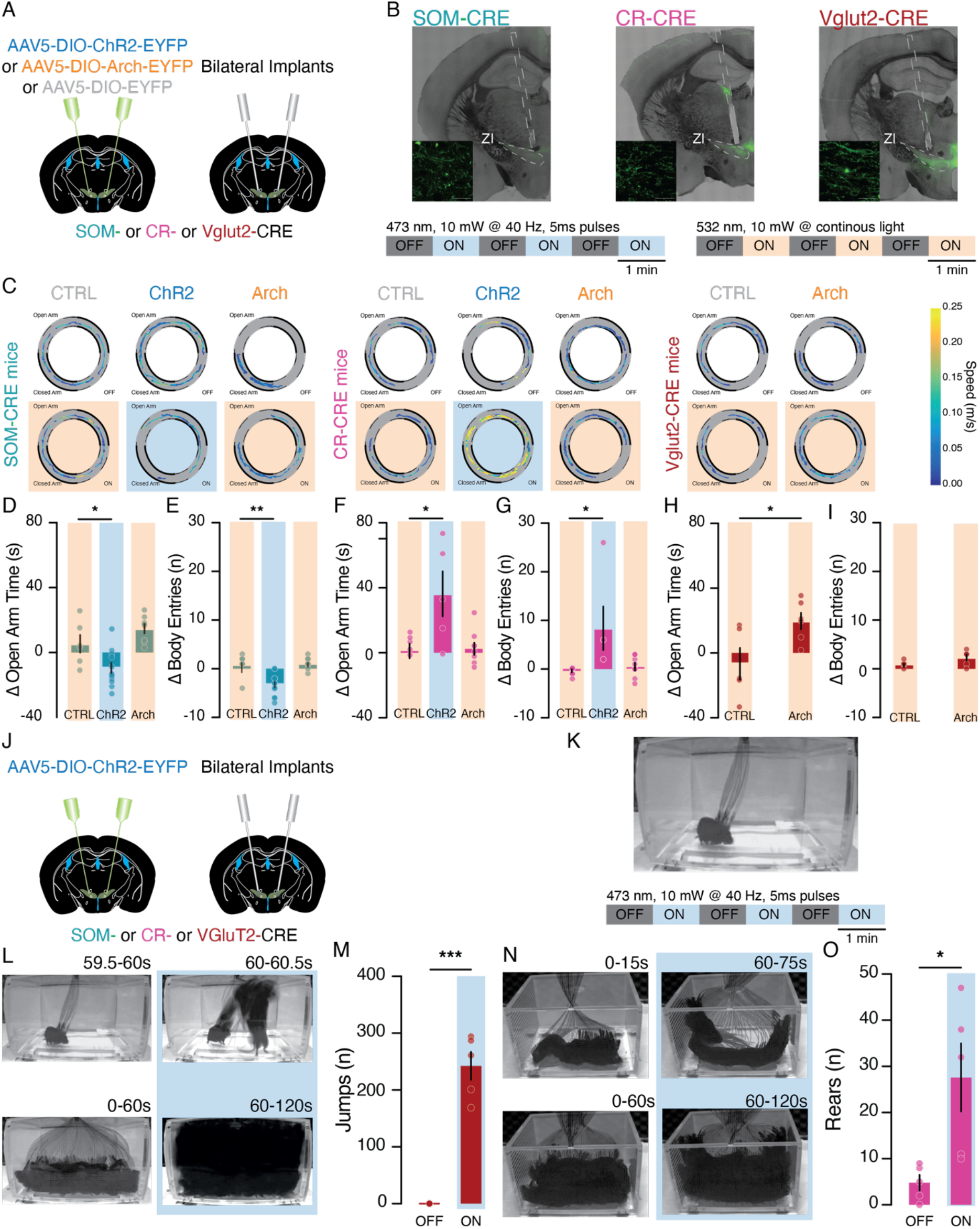
Optogenetic manipulation of SOM-, CR-, and Vglut2-expressing ZI neurons trigger distinct anxiety-related traits. A) Schematic of the experimental procedure. B) Example confocal images of coronal brain slices with the expression of ChR2 in the SOM-, CR-, and Vglut2-expressing ZI cells and the optic fiber tracts above (100μm scale bar). The inserts are high magnification zoomed-in confocal images immediately below the tip of the optic tract depicting ChR2-expressing cells (25μm scale bar). Laser stimulation protocols for ChR2 mediated activation (bottom left) and Arch mediated inhibition (bottom right) of ZI cells. C) Example track plots of the center of mice expressing control fluorescent protein, ChR2, and Arch in the SOM-, CR-, and Vglut2-expressing ZI cells on the EOM (dashed lines denote the open arms while the filled lines represents the closed arms). The periods when the laser stimulations were OFF are shown on top and ON are shown at the bottom and shaded with the speed color-coded. D) Bar graph depicting the difference in the amount of time mice expressing the control construct, ChR2, and Arch in the SOM-expressing ZI cells spent in the open arm between periods when the laser was on and off. E) Bar graph depicting the difference in the number of times mice expressing the control construct, ChR2, and Arch in the SOM-expressing ZI cells entered open arm between periods when the laser was on and off. F) Bar graph depicting the difference in the amount of time CR-CRE mice spent in the open arm between periods when the laser was on and off. G) Bar graph depicting the difference in the number of times CR-CRE mice entered open arm between periods when the laser was on and off. H) Bar graph depicting the difference in the amount of time Vglut2-CRE mice spent in the open arm between periods when the laser was on and off. I) Bar graph depicting the difference in the number of times Vglut2-CRE mice entered open arm between periods when the laser was on and off. J) Surgical setup to express excitatory opsin (ChR2) in SOM-, CR-, and Vglut2-expressing ZI cells and the implantation of optic fibers above. K) Photograph of the closed transparent chamber and the laser stimulation protocol for the ChR2 mediated activation. L) Example minimum intensity projection of a 0.5s and a 1-minute window before and after laser stimulation (blue-shaded) demonstrating the light triggered jumping behavior observed in the mice expressing ChR2 in the Vglut2-expressing ZI cells. M) Bar graph presenting the cumulative number of jumps made by mice expressing ChR2 in the Vglut2-expressing ZI cells during the laser off and on periods. N) Example minimum intensity projection of a 15s and a 1-minute window in the stimulation off and on period (blue-shaded) demonstrating the light induced increase in the rearing behavior observed in the mice expressing ChR2 in the CR-expressing ZI cells. O) Bar graph presenting the cumulative number of rears made by mice expressing ChR2 in the CR-expressing ZI cells during the laser off and on periods. are presented with the mean ± SEM.

At the behavioral level, mice performed the EOM task while the activity of SOM-, CR- and Vglut2-expressing cells was manipulated (1 min OFF, 1 min ON; 3 times, Figure 6C). Behavioral changes induced by light stimulation were reported as the difference in the performance during ON and OFF light stimulation. Activation of ChR2 expressing ZI SOM-neurons (40 Hz, 5 msec pulses) significantly reduced both the amount of time spent in the open arm (Figure 6D, Ctrl 5.40 ± 5.47 vs ChR2 −9.53 ± 3.16s, 6-12 mice) and the number of entries made into the open arm (Figure 6E, Ctrl 0.83 ± 1.01 vs ChR2 −3.26 ± 0.65 entries). In stark contrast, optogenetic activation of CR-expressing neurons induced an opposite behavioral phenotype; with a significant increase of both the time spent in the open arm (Figure 6F, Ctrl 1.98 ± 4.47 vs ChR2 36.32 ± 13.83s, 5-8 mice) and the number of open arm entries (Figure 6G, −0.67 ± 0.33 vs 8.40 ± 4.49 entries). The optogenetic inhibition of either SOM- or CR-ZI neurons had little effect on the EOM performance (Figure 6D, SOM open arm time, Ctrl 5.40 ± 5.47 vs Arch 14.74 ± 2.89s; Figure 6E, SOM open arm entry, Ctrl 0.83 ± 1.01 vs Arch 1.13 ± 0.44 entries; Figure 6F, CR open arm time, Ctrl 1.98 ± 4.47 vs Arch 3.38 ± 3.74s; Figure 6G, CR open arm entry, −0.67 ± 0.33 vs 0.50 ± 0.78 entries). Interestingly, the light-induced inactivation of Vglut2-expressing cells increased the time mice spent in the open arms (Figure 6H, Ctrl −6.64 ± 9.80 vs Arch 19.75 ± 5.14s, 5-6 mice) without affecting the number of open arm entries (Figure 6I, Ctrl 1.00 ± 0.32 vs Arch 2.33 ± 0.62 entries).

Importantly when mice were tested in the open field, the same light stimulations induced no significant change in locomotion (Figure S5) as reported by the distance travelled (Figure S5D, SOM: Ctrl −0.34 ± 0.35 vs ChR2 −1.82 ± 0.63 vs Arch 0.40 ± 0.57m; Figure S5F, CR: Ctrl −0.50 ± 0.67 vs ChR2 −0.03 ± 1.15 vs Arch 0.78 ± 0.65m; Figure S5H, Vglut2: Ctrl −0.27 ± 0.65 vs Arch 1.42 ± −0.83m) and the immobility time (Figure S5E, SOM: Ctrl −6.22 ± 6.90 vs ChR2 26.2 ± 10.29 vs Arch −2.51 ± 9.26s; Figure S5G, CR: Ctrl −7.22 ± 9.77 vs ChR2 8.28 ± 11.05 vs Arch −16.24 ± 8.11s; Figure S5I, Vglut2: Ctrl 11.18 ± 6.94 vs Arch −24.52 ± 14.03s). Photo-activation of Vglut2-expressing neurons was also tested. Due to the photo-triggered impressive jumps, testing mice in the EOM and the OFT was not possible. To measure such phenotype, mice were placed inside a closed transparent chamber (Figure 6J-K). Blue light stimulation (40Hz) triggered time locked jumps that were immediately stopped when the light stimulation was OFF (Figure 6L-M, Video S3, OFF 0.00 ± 0.00 vs ON 242.4 ± 24.60 jumps). These jumps may reflect an escape behavior.

Finally, considering that ZI CR-ChR2 expressing mice exhibited an increased exploration in the EOM, we also tested them in the closed transparent chamber. Light stimulation significantly increased the number of rears (Figure 6N-O, Video S4, OFF 4.80 ± 1.72 vs ON 27.60 ± 7.38 rears). Such exploratory rearing behavior provides an additional support that the activation of CR-expressing ZI neurons reduces anxiety. Furthermore, the activation of SOM-expressing neurons induced no clear-cut observable behavioral alterations (Video S5).

Together, our findings suggest that SOM-, CR- and Vglut2-expressing ZI subpopulations differentially modulate anxiety traits and related behaviors. Specifically, activation of SOM-expressing ZI neurons induces anxiogenesis while the activation of CR-expressing ZI neurons triggers anxiolysis and exploratory rears. Activation of Vglut2-expressing ZI neurons induces escape-like jumps and is contrasted by its inactivation, which is anxiolytic.

## Discussion

With this study, we aimed at providing experimental evidence into the mechanism by which the ZI encodes anxiety, in light of the interesting recent clinical observations reporting that ZI DBS treatment in PD reduces anxiety (*11*, *12*). Specifically, we showed that a powerful anxiety provoking stimulus such as the forced exposure on a narrow elevated platform activates ZI neurons as reported by their significant enhancement in cfos expression, adding to the growing literature that the ZI is sensitive to stressful experiences (*20*). Given the temporal limitations of such a biochemical investigation, we scrutinized the same question by examining the *in vivo* calcium dynamics of individual ZI neurons while mice explored an elevated platform. This revealed that a subset of neurons is active in the open arm edges, and become more active when mice enter a more anxiogenic area and less to a safer one. These attributes cannot be solely explained by changes in motion as most movement speed correlated ZI cells are not correlated with anxiety. Supplementing this, the local infusion of a classical anxiolytic drug, diazepam (*32*, *33*) directly into the ZI, reduced anxiety without affecting locomotion on the EOM and OFT; respectively.

It is worth noting that not all ZI cells show potentiated cfos expression or increased calcium activity following exposure to anxiogenic cues. These observations hint towards some underlying heterogeneities. Past literature provided comprehensive insights into the diversity of biochemical cell type markers expressed in the ZI (*29*–*31*), without much further characterization into their physiological and functional properties. In line with previous studies (*29*, *31*), we showed that cells expressing GABAergic markers are more prevalent over glutamatergic ones. We focused on two GABAergic subpopulations expressing Vgat with SOM or CR in addition to the rarer Vglut2-expressing cells and reported their unique intrinsic properties; leading the way with the electrophysiological profiling of genetically defined ZI subpopulations. We also provide experimental evidence that they similarly respond to anxiety-related cues biochemically and to diazepam *ex vivo*; however, they drive specific and distinct actions. Indeed, optogenetic activation of SOM-expressing cells reduced exploration in the anxiogenic open arms while the activation of CR-expressing neurons as well as inhibition of Vglut2-expressing ZI cells increased it. These findings differ from previous reports where manipulation of the ZI did not influence anxiety. Specifically, non-selective inhibition of ZI neuronal activity with either tetanus toxin or chemogenetic manipulations was reported to not significantly affect exploration in the EPM (*27*) and OFT (*34*). Moreover, chemogenetic inhibition of Vgat-expressing neurons did not reduce anxiogenic center exploration in the OFT (*34*). These differences could potentially derive from the compensatory mechanisms following permanent impairments of the ZI by the tetanus toxin in the former study, the mild nature of center in the OFT to provoke anxiety by the latter investigation, or due to differences in the specificity of cells manipulated in both cases.

In addition, we observed that the activation of Vglut2-expressing ZI cells induced time-locked jumps, characteristic of an escape-like behavior to a strong aversive stimulus (*1*, *35*), indicating the potential for Vglut2-expressing ZI cells to drive escape when the animal is faced with an anxiogenic condition. On the other hand, the activation of CR-expressing ZI cells promoted rearing, a rodent specific exploratory behavior (*36*) that is reduced in anxious animals following stressful experiences (*37*). Together with the anxiolytic effects observed in the EOM, the activation of CR-expressing ZI cells could provide the necessary foundations and partake in foraging (*18*) and hunting (*38*) behaviors, actions that have been previously attributed to the ZI. Finally, these behavioral alterations in anxiety-related behaviors that we observed with the optogenetic manipulations of its cell types are not concomitant with any significant changes in general locomotion, as is similarly reported by others (*27*, *28*, *34*).

While we bidirectionally manipulated the activity of SOM-, CR-, and Vglut2-expressing ZI neurons, the behavioral modifications were often unidirectional but still in opposition with one another. This point is best exemplified by the manipulation of SOM- and CR-expressing ZI cells, where activation of SOM- and CR-ZI cells respectively enhanced and reduced anxiety. The inability for the inhibition of these cell types to produce significant opposing modulations in anxiety may potentially be attributed to the insufficient drive by Arch to completely shut down ongoing activity for prolonged periods of time in face of strong excitatory inputs.

Together, our findings implicate that SOM-, CR-, and Vglut2-expressing ZI cells could be engaged in specific anxiety-related subcircuits. By sharing some key afferents and diazepam sensitive receptor subunits, these three ZI cell types could be similarly activated by anxiety-provoking cues and diazepam; respectively. On the other hand, they could have different outgoing projections to drive specific behavioral responses. Furthermore, biased and hierarchical local connectivity amongst these three and other ZI cell types could help explain the opposing phenotypes observed.

Building on the clinical observations, our findings provide important experimental evidence to show how the ZI and its individual subpopulations encode anxiety and guide related behaviors. Our study hence extends a positive outlook for the ZI as a therapeutic target for the treatment of anxiety.

## Materials and Methods

### Animals

Male and female SOM-CRE (B6-Sst<tm2.1(cre)Zjh>/J) (*39*), CR-CRE (B6(Cg)-Calb2<tm1(cre)Zjh>/J) (*39*), Vglut2-CRE (B6.Cg-Slc17a6<tm2(cre)Lowl>Unc/J, UNC) (*40*) and C57BL/6JRj mice were bred in-house. SOM-Tom and CR-Tom reporter lines were generated by crossing the respective Cre lines with the Ai9 transgenic line B6;129S6-Gt(ROSA)26Sor<tm9(CAG-tdTomato)Hze>/J) (*41*). BALB/cByJRj mice, previously shown to have elevated levels of anxiety (*42*, *43*), were purchased from Janvier Labs. All experimental procedures were approved by the Institutional Animal Care Office of the University of Basel and the Cantonal Veterinary Office under the License Number 2742.

### Surgery

#### General

Mice were anesthetized with gas isoflurane (5% for induction and 1.5% for maintenance) in O_2_ (Provet/Primal Healthcare, EZ Anesthesia Systems) and placed onto the stereotaxic frame (World Precision Instruments). Lidocaine (0.2mg/g) (Steuli Pharma) was injected subcutaneously above the skull. The skin was disinfected with 70% ethanol and Betadine (Mundipharma). A small incision along the anterior-posterior axis was made at the midline and then the skull was leveled. A small hole was drilled through the skull above the ZI ((*44*); 1.3mm posteriorly to the Bregma, ±1.3mm laterally to the midline, and −4.87 mm deep from the surface of the skull at a 10 degrees angle away from the midline). Glass micropipettes (Drummond) were pulled with a vertical puller (Narishige). Viral vectors were ejected into the ZI with a pressure injector system controlled by a pulse generator (A.M.P.I.; 10-100ms, 20psi, at 0.333 Hz). The wound was either stitched together with tissue absorptive silk sutures (SABANA) or continued with additional implantations (optic fiber, cannula, or gradient refractive index lens). Buprenorphine (Bupaq P, Steuli Pharama AG) pain killers were administered (0.1μg/g) post-surgery as needed.

#### Calcium Imaging

Viruses (AAV1.Camk2a.GCaMP6m.WPRE.SV40, Ready to Image virus, Inscopix) were unilaterally (150nl) injected in the ZI and then a gradient refractive lens (GRIN lens; Inscopix; 600μm wide, 7.3mm long) was placed at the same coordinates. The lens was fixed with an UV-light curable glue (Henkel). A custom-made head bar (2cm long, 0.4cm wide, 0.1cm tall) was placed for future handling. A fixed headcap was built from layers consisting of super-glue (Cyberbond), UV-light curable glue (Loctite), and dental cement (Lang). Small screws were anchored into the skull to improve adhesion between the skull and the head cap. The headcap was secured to the skin with Vetbond tissue adhesive glue (3M). The implant was covered with temporary silicone gels (KauPo Smooth-On). The expression of the GCaMP6m and the clearing of the lens were assessed regularly starting from 2 weeks post-surgery. Once the conditions were visually determined to be satisfactory, the mice were placed under isoflurane anesthesia on the stereotaxic frame and the baseplate was secured to the headcap using a UV-light curable flowable composite glue (Kerr).

#### Cannula Implantation

Custom guide cannulas (460μm in diameter and 5.5mm in length, P1 Technologies) were bilaterally placed above the ZI. The cannulas were filled with a dummy cannula of the same length and covered by temporary silicone gel (KauPo).

#### Diazepam Infusion

Following surgical recovery (> 2 weeks), internal cannulas (100μm in inner diameter and 5.5mm in length, P1 Technologies) compatible with the guide cannulas were inserted. Freshly prepared diazepam (1.5mg/kg, Fagron) in vehicle (50% propylene glycol in sterile deionized H_2_O, Sigma Aldrich) or vehicle were bilaterally infused into the ZI with the syringe pump (Harvard Apparatus) at 100nl/minute. The treatments were reversed after 2 weeks. The long gap is implemented to avoid potential acute re-exposure effects on anxiety-related behaviors in the behavioral paradigms involving elevated mazes (*45*, *46*). Following the end of all behavioral experiments, Red Retrobeads (Lumaflour) were infused via the same injection system for anatomical validation. The mice were then sacrificed and histological preparations (see section on Histology) were made.

#### Preparation of Reporter Mice for cfos Screening or in-vitro Recordings

To identify Vglut2+ ZI neurons whilst a reporter line is lacking, adult male and female (8-10 weeks) Vglut2-CRE mice were bilaterally injected with a virus expressing a CRE-dependent fluorescent protein (AAV5-EF1α-DIO-EYFP, UNC Vector Core, 300 nl/side). The virus was expressed for at least 2 weeks prior to histology or *in-vitro* electrophysiology experiments.

#### Optogenetics and Fiber Implantation

Mice were injected with viruses that contained a CRE-dependent opsin construct (AAV5-EF1α-DIO-eArch3.0-EYFP, AAV5-EF1α-DIO-ChR2-H134R-EYFP UNC/UZH Vector Cores) or a control fluorescent protein (AAV5-EF1α-DIO-EYFP, UNC Vector Core) bilaterally in the ZI (150nl/side). Optic fibers (>70% light transmittance, 200μm diameter 0.39NA optic fibers fixed by epoxy to 1.25mm wide 6.4mm long ceramic ferrules, Thorlabs, (*47*)) were bilaterally placed above the ZI (250μm).

### *In vitro* electrophysiology

Coronal mouse brain slices (180μm thick) containing the ZI were collected with a vibratome (Leica) in cooled artificial cerebrospinal fluid (ACSF) cutting solution (in mM: NMDG 92, KCl 2.5, NaH_2_PO_4_ 1.25, NaHCO_3_ 30, HEPES 20, Glucose 25, Thiourea 2, Na-ascorbate 5, Na-pyruvate 3, CaCl_2_·2H_2_O 0.5, MgSO_4_·7H_2_O 10, NAC 10; osmolarity of 290-300 mOsm) bubbled with 95% O_2_ and 5% CO_2_. The slices were kept in the holding ACSF solution (in mM: NaCl 92, KCl 2.5, NaH_2_PO_4_ 1.25, NaHCO_3_ 30, HEPES 20, Glucose 25, Thiourea 2, Na-ascorbate 5, Na-pyruvate 3, CaCl_2_·2H_2_O 2, MgSO_4_·7H_2_O 2, NAC 10; pH 7.3; osmolarity of 290-300 mOsm) at 31 °C before transferred into the recording chamber containing the recording ACSF solution (in mM: NaCl 119, KCl 2.5, NaH_2_PO_4_ 1.25, NaHCO_3_ 24, Glucose 12.5, CaCl_2_·2H_2_O 2, MgSO_4_·7H_2_O 2; pH 7.3; osmolarity of 290-300 mOsm) bubbled with 95% O_2_ and 5% CO_2_. Visualized whole-cell patch recordings were performed to record cells from the ZI. We identified SOM-, CR-, and Vglut2-positive cells by the expression of TdTomato from reporter lines or virally introduced fluorescent proteins (AAV5-EF1α-DIO-EYFP, AAV5-EF1α-DIO-eArch3.0-EYFP, AAV5-EF1α-DIO-ChR2-H134R-EYFP) in each of the three cell-type specific CRE lines (SOM-CRE, CR-CRE, and Vglut2-CRE) visualized under an upright microscope (OLYMPUS) through PLANFL N 4X and 40X (OLYMPUS) objectives. The internal solution contained; for electrophysiological profiling and optogenetic tools validation (in mM: K-Gluconate 130, Creatine Phosphate 10, MgCl_2_ 4, Na_2_ATP 3.4, Na_3_GTP 0.1, EGTA 1.1, HEPES 5; pH 7.3; osmolarity of 289 mOsm) and for IPSC recordings (in mM: K-Gluconate 30, KCl 100, Creatine Phosphate 10, MgCl_2_ 4, Na_2_ATP 3.4, Na_3_GTP 0.1, EGTA 1.1, HEPES 5; pH 7.3; osmolarity of 289 mOsm).

To investigate the ex vivo effect of diazepam, diazepam (1μM, Fagron) was subsequently washed in and blocked with picrotoxin (100μM). In separate slices, spontaneous inhibitory post-synaptic potentials were recorded and also subsequently bath applied with diazepam and blocked with picrotoxin. For the validation of optogenetic tools, light flashes (400ms) controlled by a pulse generator (A.M.P.I) were delivered from a microscope mounted LED (CoolLED) to evoke synaptic currents in voltage clamp mode held at resting membrane potential. In current clamp mode, blue light flashes (1s continuous and 10s at 40Hz) were used to evoke firing in ChR2-expressing cells and orange light flashes (1s continuous) were used to reduce electrically evoked (current injected) firing.

### Histology

#### Immunohistochemistry

Mice received a lethal dose of pentobarbital (0.3mg/kg i.p.) (Steuli Pharama AG). The animals were then transcardially perfused with 25mL of phosphate buffered saline buffer (PBS) (Sigma Aldrich) followed by the same volume of 4% paraformaldehyde (PFA) (Sigma Aldrich) in PBS. Brains were extracted and post-fixed in 4% PFA overnight then immersed in 30% sucrose in PBS until the brains has completely sunk to the floor of the container (complete absorption of sucrose). Sections were prepared (50μm) using a cryostat (Leica 1950 CM). Brain slices containing the ZI slices were first washed (3×3 mins) with 1X Tris-buffered saline (TBS) (Sigma Aldrich) with 0.1% Tween-20 (Sigma Aldrich), permeabilized for 20mins with 1X TBS with 0.1% Tween-20 and Triton-X100 (Sigma Aldrich) and blocked for 120 mins with 1% Bovine Serum Albumin (Sigma Aldrich) or 1% Bovine Serum Albumin with 10% Donkey Serum (Sigma Aldrich) in 1X TBS with 0.1% Tween-20. The brain slices were stained overnight in the blocking solution with the primary antibody (Rabbit anti-cfos, abcam ab7963, 1:500 and/or Goat anti-CR, swant CG1, 1:1000). In the following day, the brain slices were first washed and then stained for 120 minutes with the secondary antibody (Goat anti-Rabbit conjugated to AlexaFlour555, Invitrogen A21429, 1:500, Donkey anti-Rabbit conjugated to AlexaFlour488, Invitrogen A21206 1:500, or Donkey anti-Goat conjugated to AlexaFlour647, abcam ab150131, 1:500). The brain slices were again washed and then mounted on glass slides (Superfrost Plus, Thermo Scientific), immersed by DAPI containing ProLong Gold Antifade mounting solution (Invitrogen) and covered with borosilicate cover glass (VWR).

#### Fluorescent in-situ hybridization (FISH)

Mice were anesthetized with gas isoflurane. The brains were extracted and quickly frozen on aluminum foil over dry ice. Brains were kept overnight at −80°C. On the next day, the brains were equilibrated at −20°C in the cryostat for at least 30 minutes. Fresh frozen brain sections (20μm) containing the ZI were prepared and directly mounted onto glass slides. Selected slices were fixed for 30 minutes using 4% PFA in PBS, dehydrated with increasing concentrations of ethanol (50%, 70%, 100%, 100%), surrounded by hydrophobic barriers (ImmEdge, Vector Laboratories), before proceeding with the RNAscope Fluorescent Multiplex (ACD, Biotechne) Assay to label the mRNAs for vesicular GABA transporter (Vgat, Slc32a1, ref319191), vesicular glutamate transporter-2 (Vglut2, Slc17a6, ref319171), somatostatin (SOM, SST, ref482691), calretinin (CR, CALB2, ref313641), and parvalbumin (PV, Pvalb, ref421931). Processed slides were immersed in ProLong Diamond Antifade mounting solution (Invitrogen) and covered with borosilicate cover glass.

#### Confocal Imaging

To visualize fluorescent IHC, FISH, or reporter protein labeling in brain sections, mounted sections were imaged with a Zeiss LSM700 upright confocal microscope controlled by the ZEN Black image acquisition software (v2010, Zeiss). PLAN APO 20X (0.8NA, air) and 40X (1.3NA, oil) objectives were used to take low and high magnification images, respectively. Fixed wavelength (405nm, 488nm, 555nm, and 639nm) lasers were utilized to visualize specific fluorescent signals while brightfield imaging was used for the anatomical features. Images were stitched with the acquisition software and further post-processed in FIJI (v2.0.0, ImageJ).

### Behavior

#### Elevated Rod Exposure for cfos Screening

Mice were forcibly confined on an elevated rod (100cm tall, 3cm diameter) for 30 minutes and put back into the home cage. Soft packing foam was placed around the bottom of the elevated rod. Control mice were left in the home cage for the same duration. Mice were transcardially perfused 30 minutes later.

#### Elevated Platform Exposure for in-vivo Calcium Imaging

Mice were connected to the nVoke2 (Inscopix) system and recorded with the Inscopix data acquisition software with a 20Hz sampling rate while behaviorally recorded simultaneously from above using a camera (The Imaging Source) that is controlled by ANYMAZE (v5.23, Stoeling) and sampled up to 20Hz. The mice were then placed in the enclosed center of the elevated platform and recorded while freely moving for 5 minutes. The elevated platform is constructed by completely blocking the entrance of the closed arms of an elevated plus maze (arms with 7.5cm in width, 30cm in length separated by a center that is 7.5cm in width and in length and enclosed by 60cm tall cardboard pieces on the two sides not connected to the arms). This elevated platform maximizes and forces the recorded mice to explore the anxiogenic exposed open edges. The larger dimensions of this platform also provided more room for the recorded mice to maneuver.

#### Optogenetic Stimulation

The light power for optogenetic stimulation (473nm or 532nm) were adjusted with a power-meter (Thorlabs) such that the illumination at the tip of the optic fibers would be at around 10mW. To sufficiently activate ZI neurons with ChR2, 5msec pulses at 40Hz (473nm). Constant light (532nm) stimulation were used to drive the Arch3.0 mediated inhibition. The number of light stimulation periods were counterbalanced with periods where no light was provided in an alternating manner over 6 minutes with 1-minute bins.

#### Elevated O-Maze (EOM)

Cannula implanted mice that received intracranial infusion of either diazepam or vehicle (see Surgery/Diazepam Infusion) or fiber implanted mice that were attached to the laser cables (see Behavior/Optogenetic Manipulations) were placed on the open arm of the EOM (arms with 5cm in width; the maze is 55cm in diameter and 60cm above the ground) facing towards the closed arm (enclosed by two 15cm tall grey opaque plastic walls). The translucent floor allowed a camera mounted below facing upwards to record the shadow of the tested mice as it traveled along the EOM. Videos were recorded and tracked with ANYMAZE at a rate of up to 30Hz for either 10 minutes (infusion animals) or 6 minutes (optogenetics animals).

#### Open Field Test (OFT)

Mice were placed in the center of a rectangular OFT box (30cm in width and 45 cm in length, with 30cm tall walls) for either 10 minutes (infusion animals) or 6 minutes (optogenetics animals). Videos were recorded with a downward-facing camera from above with ANYMAZE in the same manner as mentioned previously.

#### Unique Behaviors in the Closed Chamber

Mice were placed in the center of a transparent plexi-glass rectangular box (25cm in width and 20cm in length, with 15cm tall walls) with a closed ceiling that had a circular (5cm in diameter) hole that let out the fiber cables but not the mice for 6 minutes. Videos were recorded with a front-facing camera from the side with ANYMAZE in the same manner as mentioned previously.

### Analysis

#### Calcium Imaging

The position of the center of the video recorded mice and its movement speed were extracted from ANYMAZE together with the digital boundaries of the center and the open arms of the elevated platform. The behavioral trace was then exported to Matlab (v2018b, Mathworks) for further analysis.

Calcium transients recorded from the nVoke2 system were pre-processed in the Inscopix Data Processing Software (IDPS, v1.3, Inscopix). The time-stamped calcium traces were also exported to Matlab. Calcium transients and the behavioral traces were first aligned to each other. Calcium traces were denoised and standardized. Calcium and behavior traces were then both binned to 500ms windows. To examine the relationship between anxiety and the neuronal activity in the ZI, correlational analyses were made between the distance of the recorded mice to the enclosed center (center distance) and the calcium transients of the recorded cells. Center distance is an assessment for anxiety because the further the animal is from the center, the more exposed the animal is and thus the higher the level of anxiety. The center distance is calculated by subtracting the lengthwise position of the center of animal from that of the center of the platform and taking the absolute value. Pearson’s correlation coefficient was calculated from the distribution of the binned calcium activity and center distance values. To characterize cells as being positively, negatively, or not correlated between these two parameters, a bootstrap test was performed by randomly shifting the calcium traces 1000 times in a circular manner and then calculating the correlation coefficient. Cells whose real correlation coefficient was negative and below the 5^th^ percentile or positive and above the 95^th^ percentile of the random correlation coefficients were classified as negatively or positively correlated, respectively. Cells that did not fall into this range were classified as not correlated.

To further show that these correlated cells displayed activity with respect to the level of anxiety that the recorded mice experienced, the difference in activity between the time when the animals was in the center and the edges (defined as an area on extremities of the open arm with the same size as the center) was calculated for each cell and compared across groups. All recorded animals spent more time in the center as compared to the edges. To reduce sampling bias, the activity in the center was randomly sampled 1000 times to match the number of bins the animals spent in the edges and the averaged difference was reported.

To examine whether the correlated cells displayed changes in activity as the recorded animals transitioned between areas that is less anxiogenic to more anxiogenic and vice versa, calcium transients surrounding (2.5s before and after) the entry or the exit of the edge or the center were extracted, grouped, and compared. To visualize changes across cells and groups, transients were adjusted such that the average activity in the baseline window before the change in compartment is 0.

To show that the center distance correlated cells encoded anxiety rather than movement speed, the same correlation analysis was applied between calcium transients and movement speed for each cell. For each categorized (positively, negatively, and not correlated) center distance correlated cell groups, the proportion each of the three categories of movement speed correlated cells were shown.

#### Behavioral Paradigms

All parameters were extracted from ANYMAZE (EOM and OFT). Unique behaviors such as jumps and rears were visually identified and manually quantified.

#### Statistics

All statistical comparisons, unless otherwise noted, were analyzed with Prism (v8.4.3, GraphPad). The significance level was set at 5% (α = 0.05) in all cases. Student’s t-test was used to make comparisons between two groups of data. The unpaired design was utilized when the samples were independent from one another (number of cfos labelled ZI cells prepared from mice that were exposed to the elevated anxious experience or the control home conditions) while the paired design was applied in cases where repeated measures were made (activity of center distance cells before and after compartment change, effect of diazepam or vehicle on the performance in the EOM and OFT of the same mice, effect of diazepam on the recorded inhibitory currents onto ZI cells with respect to the baseline period before, and effect of photo-activation of CR- and Vglut2-ZI cells to respectively trigger rears and jumps as compared to periods in the same session where the light were off). One-way analysis of variance (ANOVA) with Bonferroni’s multiple comparison tests were used to compare the differences across multiple (greater than two) independent samples (differences in activity between the center and the edge across the different center distance correlated groups, electrophysiological properties among the SOM-, CR-, and Vglut2-expressing ZI cells, and the differences in behavioral parameters among control, ChR2-, and Arch-expressing groups).

## General

We thank Tania Barkat-Rinaldi and Jan Gruendemann as well as the past and current members of the Tan Lab for constructive input; the Biozentrum Imaging Facility for microscopy training, and the Inscopix team particularly Diane Damez-Werno for technical support of the calcium imaging experiment.

## Author contributions

Z. Li, G. Rizzi, and K.R. Tan all contributed to the experimental design. G. Rizzi conducted the calcium imaging experiments. K.R. Tan conducted the electrophysiology experiments. Z. Li conducted all other experiments. Z. Li and K.R. Tan analyzed the data and prepared the manuscript.

## Data and materials availability

The datasets generated during and/or analyzed during the current study as well as the custom written MATLAB scripts are available from the corresponding author upon reasonable request.

## Supplementary Materials

**Figure S1.**
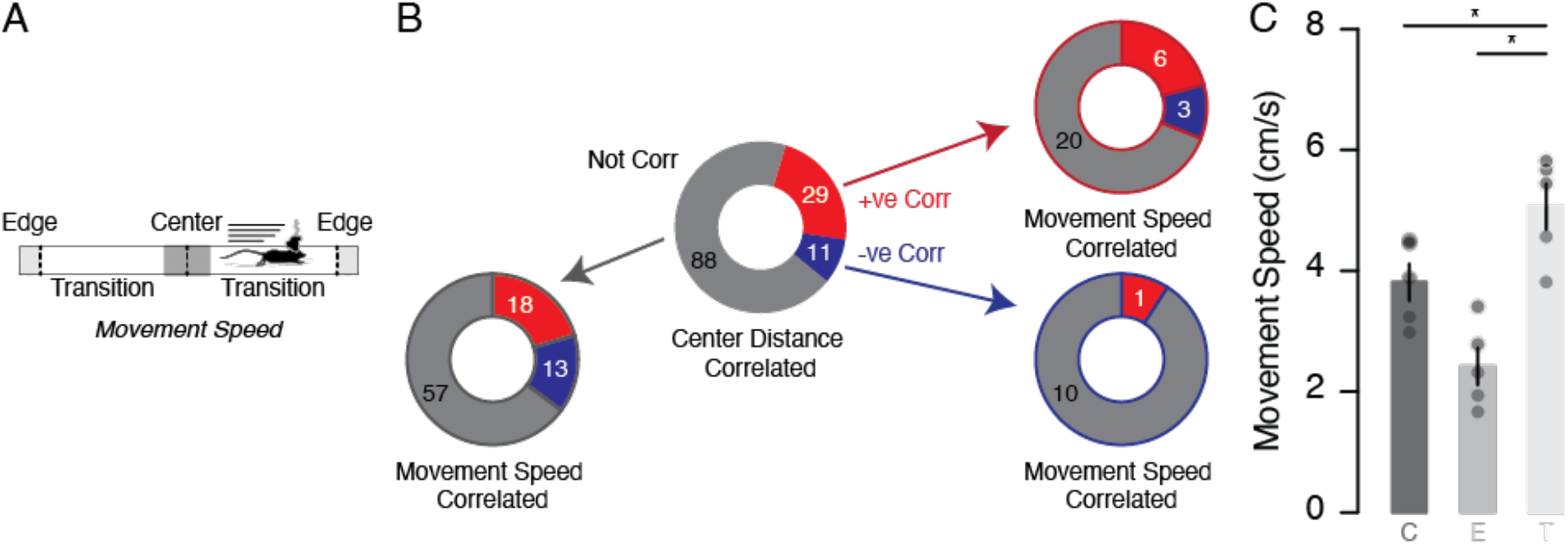
Anxiety sensitive ZI neurons are not sensitive to movement. A) Schematic of the top view of the elevated platform. B) Pie chart depicting the proportion of center distance correlated cells (center; also reported in figure 2H). Proportion of positively (lined in red, top right), negatively (lined in blue, bottom left) and not (lined in black, bottom left) correlated center distance cell groups that show movement speed correlated activity. C) Bar graphs reporting the average movement speed of the recorded mice in the center, the edges, and the transition areas. Data are presented with the mean ± SEM.

**Figure S2.**
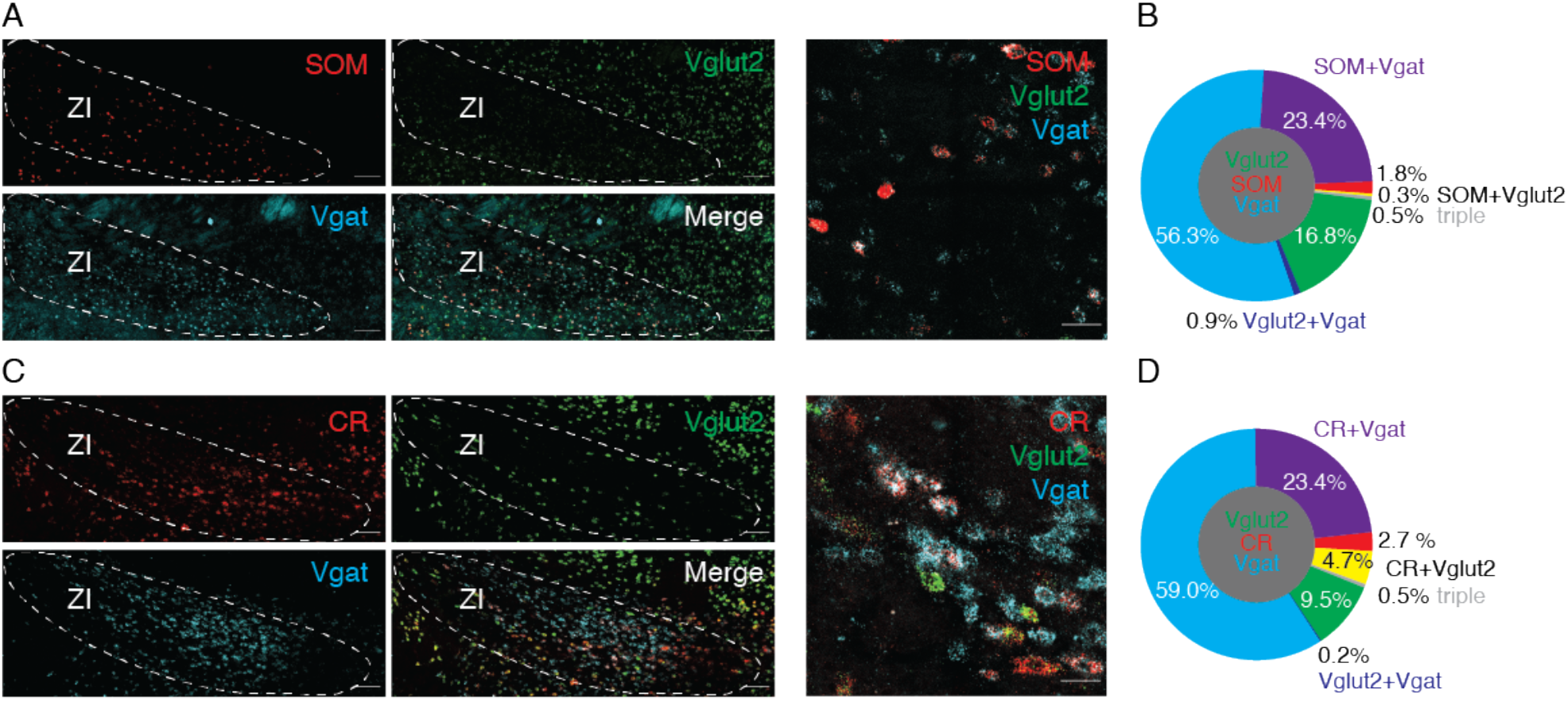
SOM- and CR-expressing ZI neurons co-express Vgat. A) Example confocal images of the ZI- containing brain slices with the FISH labeling of the mRNAs for SOM (red), Vglut2 (green), Vgat (cyan) as well as the merged image (100μm and 25μm scale bar respectively). B) Pie chart quantification of the proportion of cells in ZI expressing SOM, Vglut2, Vgat, or any combination of all three (3686 cells, 3 mice). C) Example confocal images of the ZI with the FISH labeling of the mRNAs for CR (red), Vglut2 (green), Vgat (cyan), and the merged image (100μm and 25μm scale bar respectively). D) Pie chart representation of percentage of cells in the ZI with the labeling for CR, Vglut2, Vgat, or any combination of the three (6607 cells, 3 mice).

**Figure S3.**
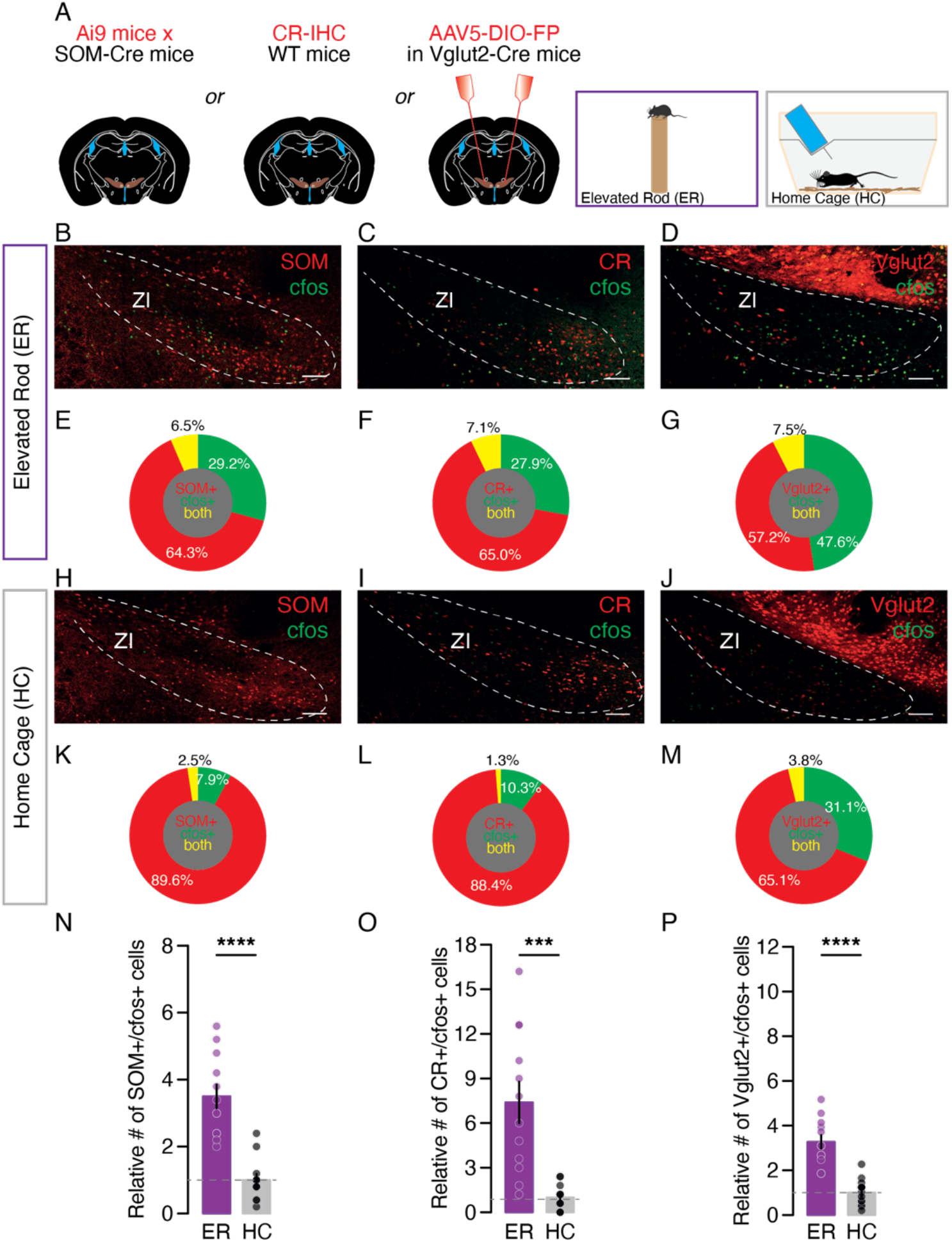
Anxious elevated experience activates ZI SOM-, CR-, and Vglut2-expressing cells. A) Schematic of the experimental procedure. B) Example confocal image of the ZI-containing brain section with SOM (red) and IHC labeling of cfos (green) in mice that were exposed to the anxious elevated rod experience (100μm scale bar). C) Example confocal image of the ZI-containing brain slice with the IHC labeling of CR (red) and cfos (green) in mice that were exposed to the anxious elevated rod experience (100μm scale bar). D) Example confocal image of the ZI with Vglut2 (red) and IHC labeling of cfos (green) in mice that were exposed to the anxious elevated rod experience (100μm scale bar). E) Pie chart reporting the proportion of ZI cells (Figure S3B) labeled with SOM, cfos, or both (3262 cells, 3 mice) F) Pie chart depicting the proportion of ZI cells (Figure S3C) labeled with CR, cfos, or both (2074 cells, 3 mice). G) Pie chart of the proportion of ZI cells (Figure S3D) labeled with Vglut2, cfos, or both (2538 cells, 3 mice). H) Example confocal image of the ZI with SOM (red) and the IHC labeling of cfos (green) in the ZI of mice that were left in the home cage (100μm scale bar). I) Example confocal image of ZI-containing brain section with the IHC labeling against CR (red) and cfos (green) in mice that were left in the home cage (100μm scale bar). J) Example confocal image of the ZI-containing brain slice with Vglut2 (red) and IHC labeling for cfos (green) in the mice that were left in the home cage (100μm scale bar). E) Pie chart representation of the proportion of ZI cells (Figure S3H) labeled with SOM, cfos, or both (2458 cells, 3 mice). F) Pie chart description of the proportion of ZI cells (Figure S3I) labeled with CR, cfos, or both (1624 cells, 3 mice); G) Pie chart quantification of the proportion of ZI cells (Figure S3J) labeled with Vglut2, cfos, or both (1551 cells, 3 mice). N) Bar graphs reporting the relative number of SOM+ and cfos+ labeled cells (normalized to the control mice) in the ZI for both experimental conditions. O) Bar graphs presenting the relative number of CR+ and cfos+ labeled cells (normalized to the control mice) in the ZI for both experimental conditions. N) Bar graphs comparing the relative number of Vglut2+ and cfos+ labeled cells (normalized to the control mice) in the ZI for both experimental groups. Data are presented with the mean ± SEM.

**Figure S4.**
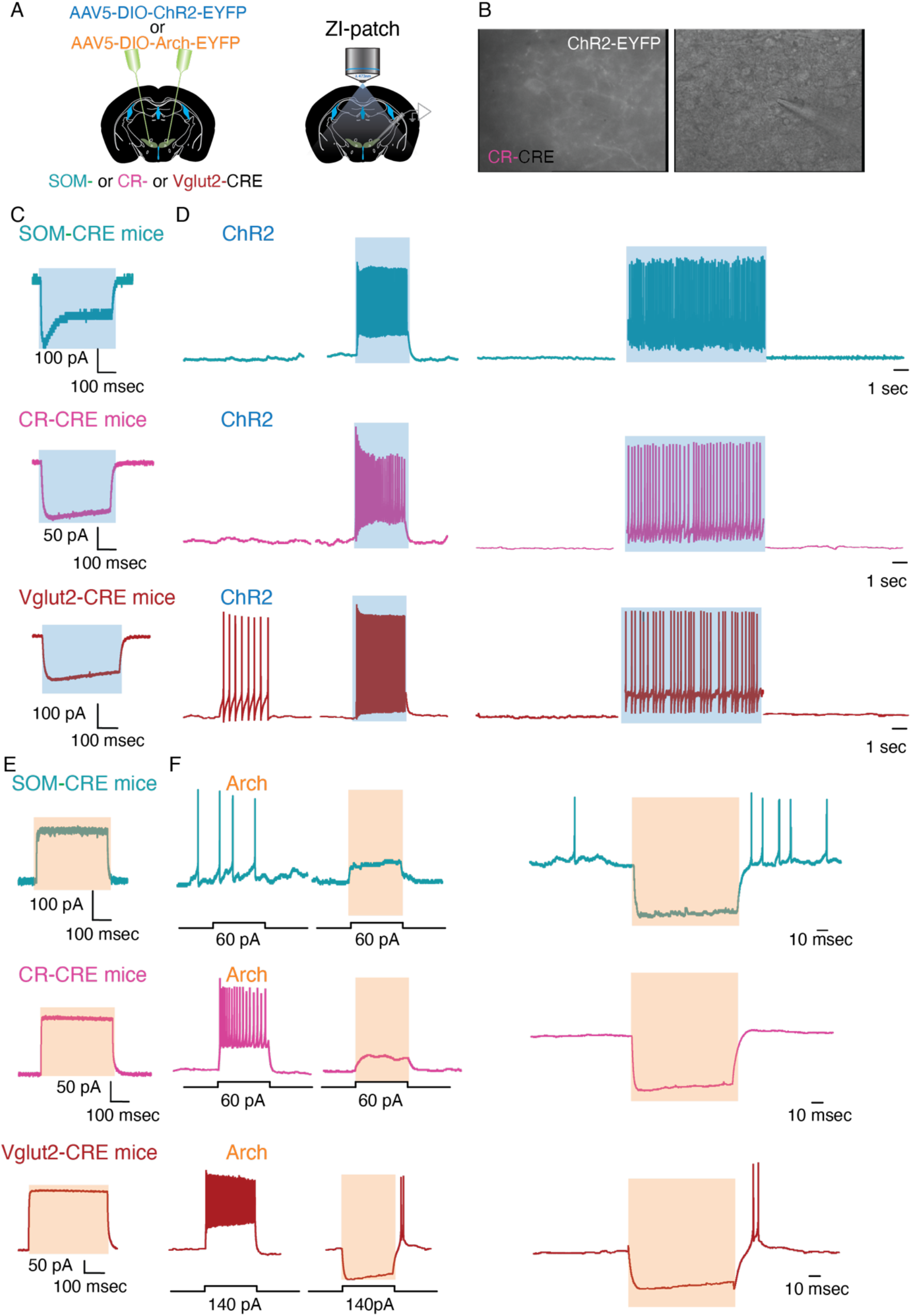
Validation of optogenetic tools in SOM-, CR-, and Vglut2-expressing ZI *in-vitro*. A) Schematic of the experimental procedure. B) Example fluorescent and brightfield images of a patched ChR2 expressing ZI CR cell. C) Example photocurrent recorded upon blue light (473nm; 400ms) illumination in representative ChR2 infected SOM- (cyan), CR- (magenta), and Vglut2- (maroon) expressing ZI cells. D) Example traces of representative ChR2 infected SOM- (cyan), CR- (magenta), and Vglut2- (maroon) expressing ZI cells recorded in current clamp mode (I = 0) that shows a continuous blue light stimulation (1s) evokes time-locked increase in firing and a 10s blue light stimulation (5ms pulses, 40Hz) also induced sustained firing in all cells. E) Example photocurrent recorded upon green light (532nm; 400ms) illumination in a representative Arch infected SOM- (cyan), CR- (magenta), and Vglut2- (maroon) expressing ZI cell. F) Example traces of a representative Arch infected SOM- (cyan), CR- (magenta), and Vglut2- (maroon) expressing ZI cell recorded in current clamp mode that depicts a continuous green light stimulation (1s) suppresses evoked firing and hyperpolarizes neurons at rest.

**Figure S5.**
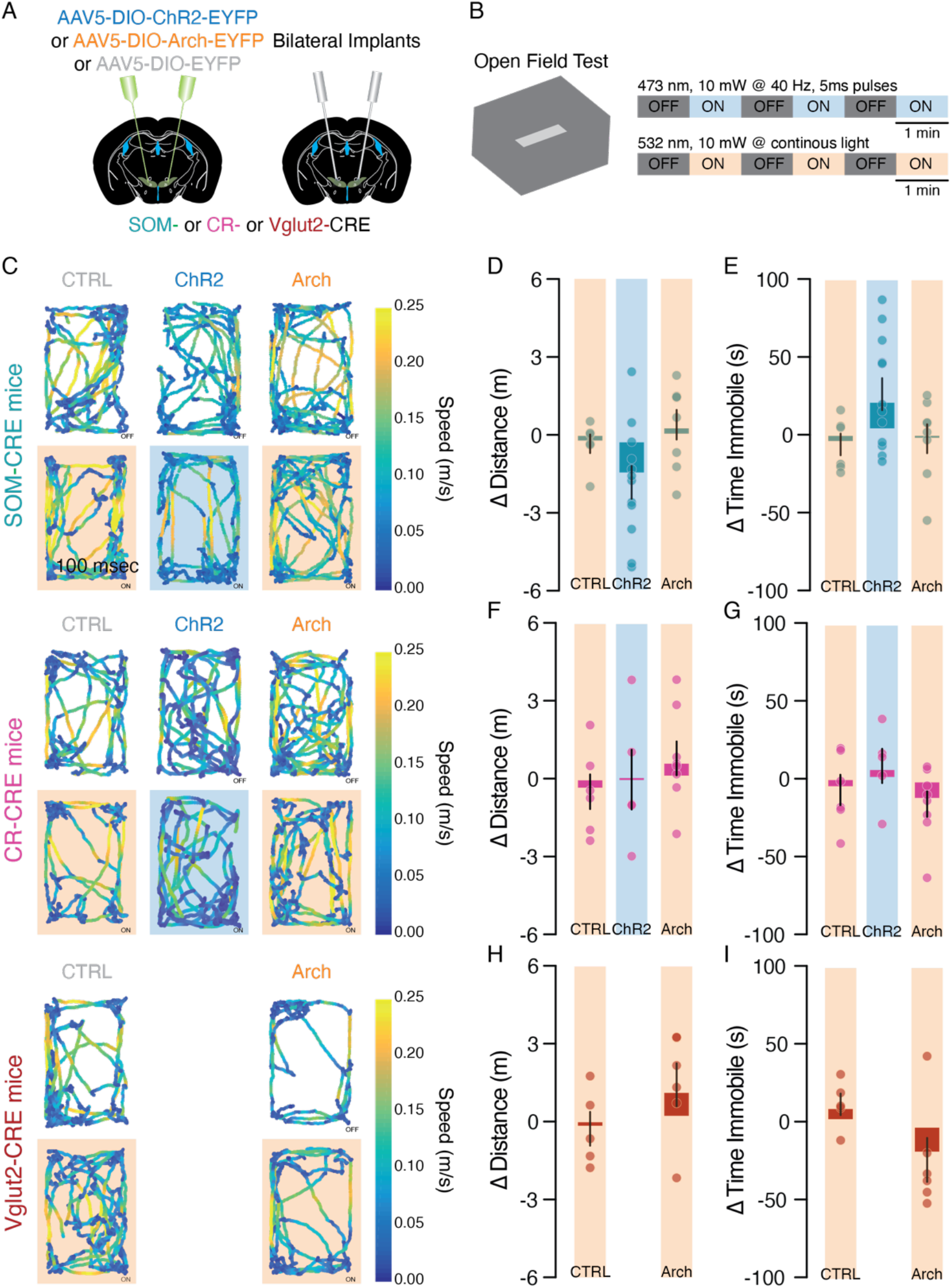
Bidirectional optogenetic manipulations of SOM-, CR-, and Vglut2- ZI cells do not impact general locomotion. A) Schematic of the experimental procedure. B) Schematic of the open field test (OFT) box and the laser stimulation protocols for ChR2 mediated activation and Arch mediated inhibition. C) Example track plots of the center of mice expressing control fluorescent protein, ChR2, and Arch in the SOM- (cyan), CR- (magenta), and Vglut2- (maroon) expressing ZI cells in the OFT during the periods when the laser stimulations were off and on (shaded) with the speed color-coded. D) Bar graph reporting the difference in distance mice expressing the control construct, ChR2, and Arch in the SOM-expressing ZI cells traveled between periods when the laser was on and off. E) Bar graph depicting the difference in the time mice expressing the control construct, ChR2, and Arch in the SOM-expressing ZI cells spent immobile between periods when the laser was on and off. F) Bar graph presenting the difference in the distance mice expressing the control construct, ChR2, and Arch in the CR-expressing ZI cells traveled between periods when the laser was on and off. G) Bar graph of the difference in the time mice expressing the control construct, ChR2, and Arch in the CR- expressing ZI cells spent immobile between periods when the laser was on and off. H) Bar graph comparing the difference in the distance mice expressing the control construct and Arch in the Vglut2-expressing ZI cells traveled between periods when the laser was on and off. I) Bar graph with the difference in the time mice expressing the control construct and Arch in the Vglut2-expressing ZI cells spent immobile between periods when the laser was on and off. Data are presented with the mean ± SEM.

**Video S1. Example calcium recording of ZI cells while the recorded mouse traveled between the enclosed center and the open edge of elevated platform.**

**Video S2. Local ZI infusion of diazepam increases open arm exploration in the EOM (4X speed).**

**Video S3. Optogenetic activation of Vglut2- ZI cells trigger time-locked jumps (4X speed).**

**Video S4. Optogenetic activation of CR- ZI cells promote rears (4X speed).**

**Video S5. Optogenetic activation of SOM- ZI cells do not elicit any peculiar behaviors (4X speed).**

**Table S1.**
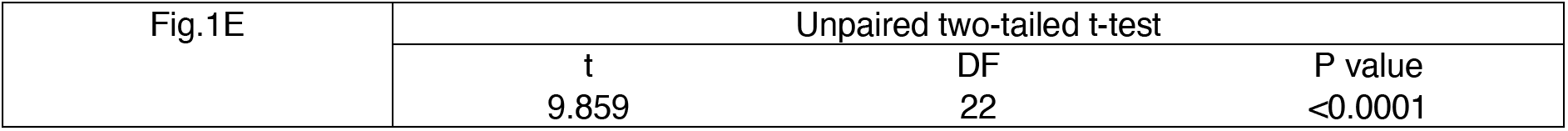
Statistical Information for Figure 1.

**Table S2.**
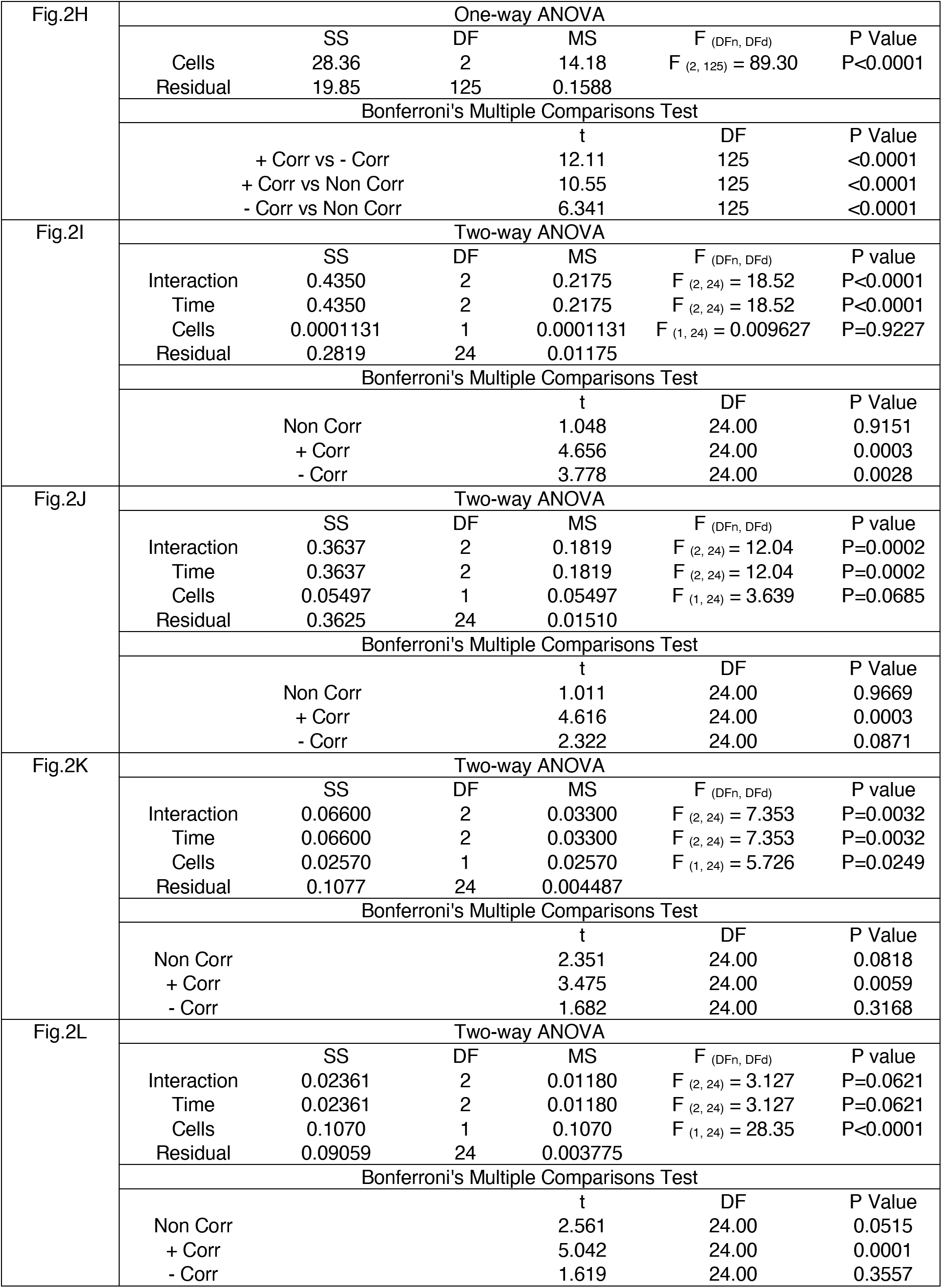
Statistical Information for Figure 2.

**Table S3.** Statistical Information for Figure 3.

|  |  |  |  |
| --- | --- | --- | --- |
| Fig.3E | Paired two-tailed t-test |  |  |
|  | t | DF | P value |
|  | 2.608 | 12 | 0.0229 |
| Fig.3F | Paired two-tailed t-test |  |  |
|  | t | DF | P value |
|  | 2.350 | 12 | 0.0367 |
| Fig.3H | Paired two-tailed t-test |  |  |
|  | t | DF | P value |
|  | 0.8318 | 12 | 0.4316 |
| Fig.3I | Paired two-tailed t-test |  |  |
|  | t | DF | P value |
|  | 0.5914 | 12 | 0.5652 |

**Table S4.**
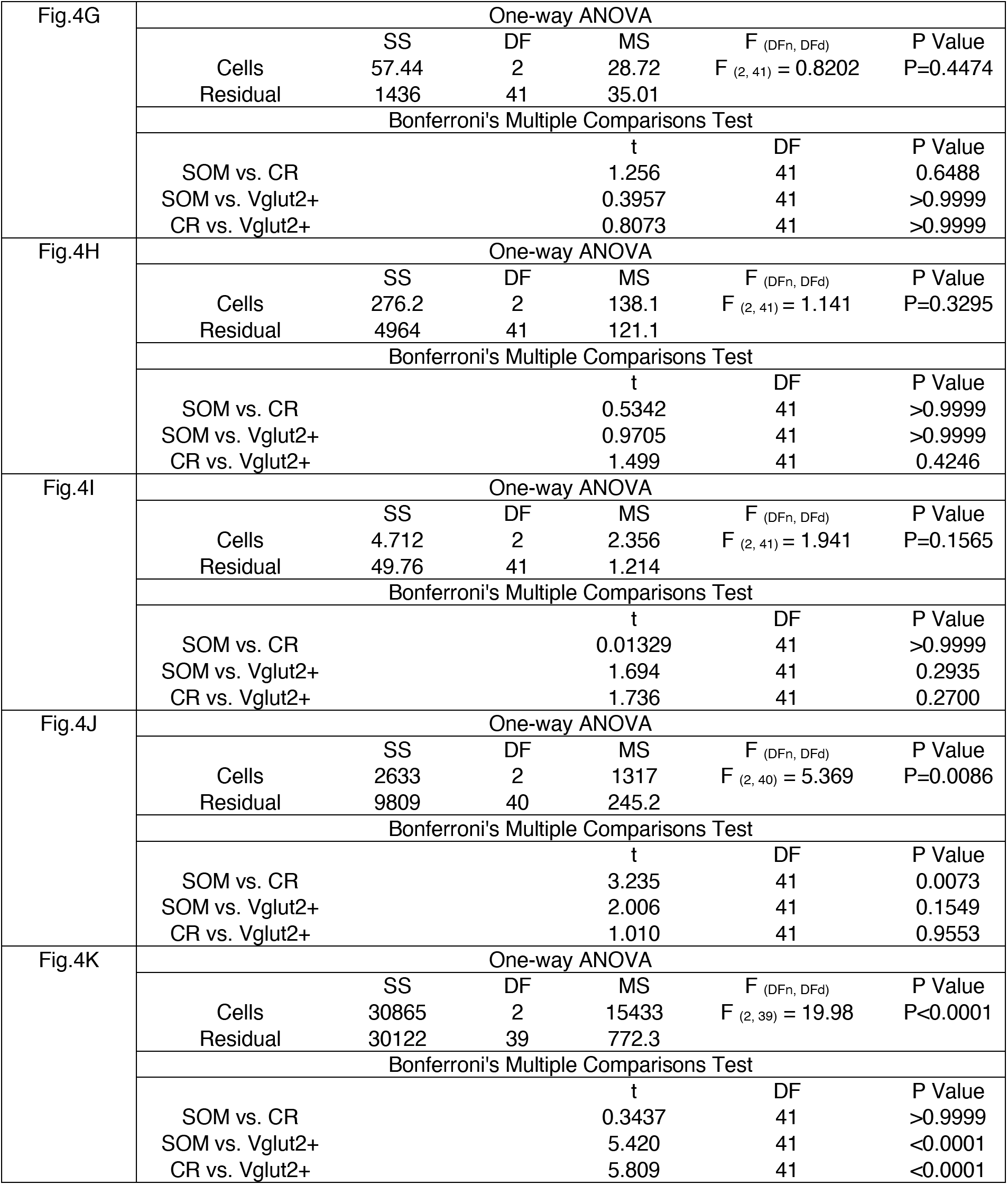
Statistical Information for Figure 4.

**Table S5.**
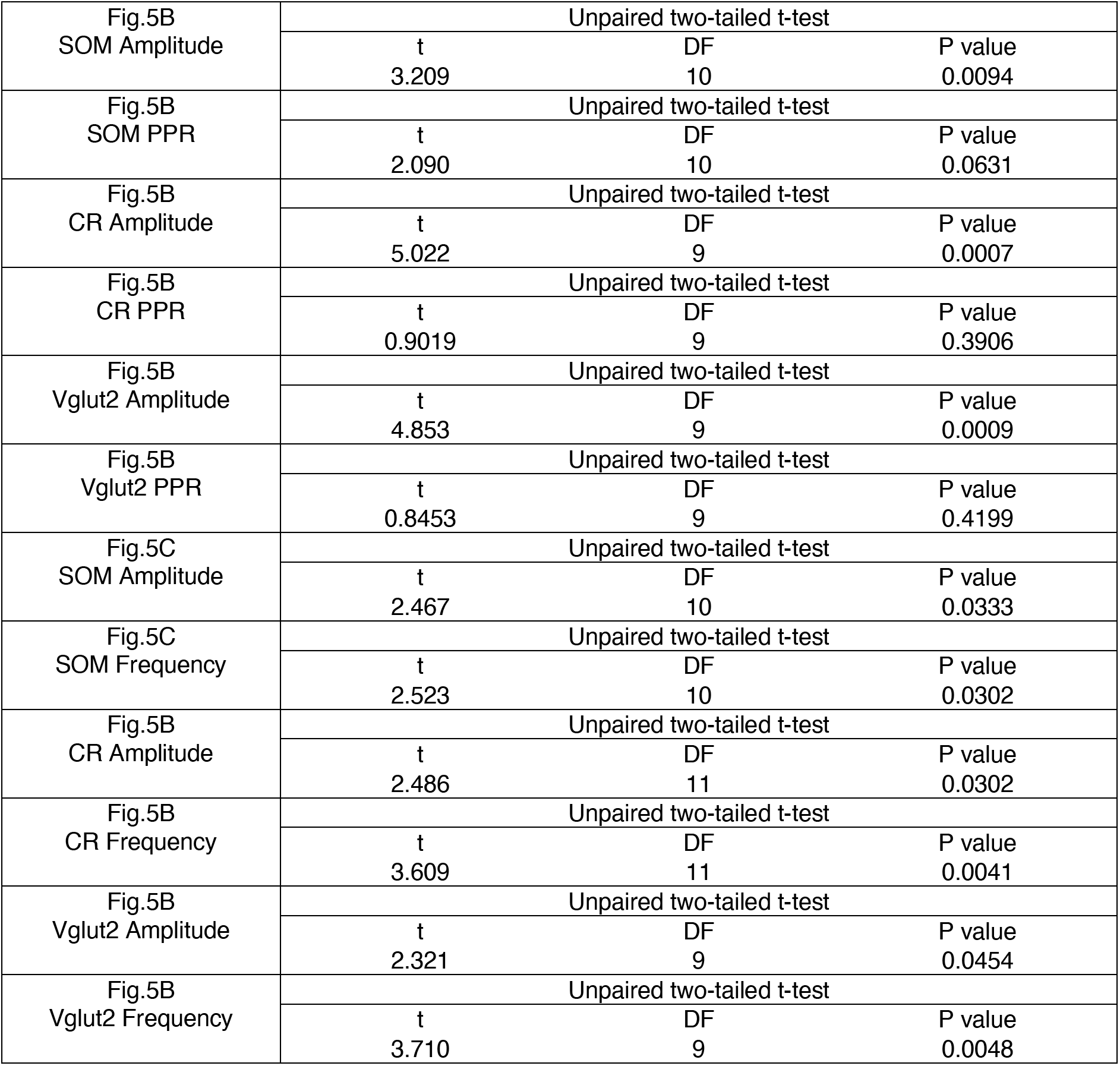
Statistical Information for Figure 5.

|  |  |  |  |
| --- | --- | --- | --- |
| Fig.5B<br>SOM Amplitude | Unpaired two-tailed t-test |  |  |
|  | t | DF | P value |
|  | 3.209 | 10 | 0.0094 |
| Fig.5B<br>SOM PPR | Unpaired two-tailed t-test |  |  |
|  | t | DF | P value |
|  | 2.090 | 10 | 0.0631 |
| Fig.5B<br>CR Amplitude | Unpaired two-tailed t-test |  |  |
|  | t | DF | P value |
|  | 5.022 | 9 | 0.0007 |
| Fig.5B<br>CR PPR | Unpaired two-tailed t-test |  |  |
|  | t | DF | P value |
|  | 0.9019 | 9 | 0.3906 |
| Fig.5B<br>Vglut2 Amplitude | Unpaired two-tailed t-test |  |  |
|  | t | DF | P value |
|  | 4.853 | 9 | 0.0009 |
| Fig.5B<br>Vglut2 PPR | Unpaired two-tailed t-test |  |  |
|  | t | DF | P value |
|  | 0.8453 | 9 | 0.4199 |
| Fig.5C<br>SOM Amplitude | Unpaired two-tailed t-test |  |  |
|  | t | DF | P value |
|  | 2.467 | 10 | 0.0333 |
| Fig.5C<br>SOM Frequency | Unpaired two-tailed t-test |  |  |
|  | t | DF | P value |
|  | 2.523 | 10 | 0.0302 |
| Fig.5B<br>CR Amplitude | Unpaired two-tailed t-test |  |  |
|  | t | DF | P value |
|  | 2.486 | 11 | 0.0302 |
| Fig.5B<br>CR Frequency | Unpaired two-tailed t-test |  |  |
|  | t | DF | P value |
|  | 3.609 | 11 | 0.0041 |
| Fig.5B<br>Vglut2 Amplitude | Unpaired two-tailed t-test |  |  |
|  | t | DF | P value |
|  | 2.321 | 9 | 0.0454 |
| Fig.5B<br>Vglut2 Frequency | Unpaired two-tailed t-test |  |  |
|  | t | DF | P value |
|  | 3.710 | 9 | 0.0048 |

**Table S6.**
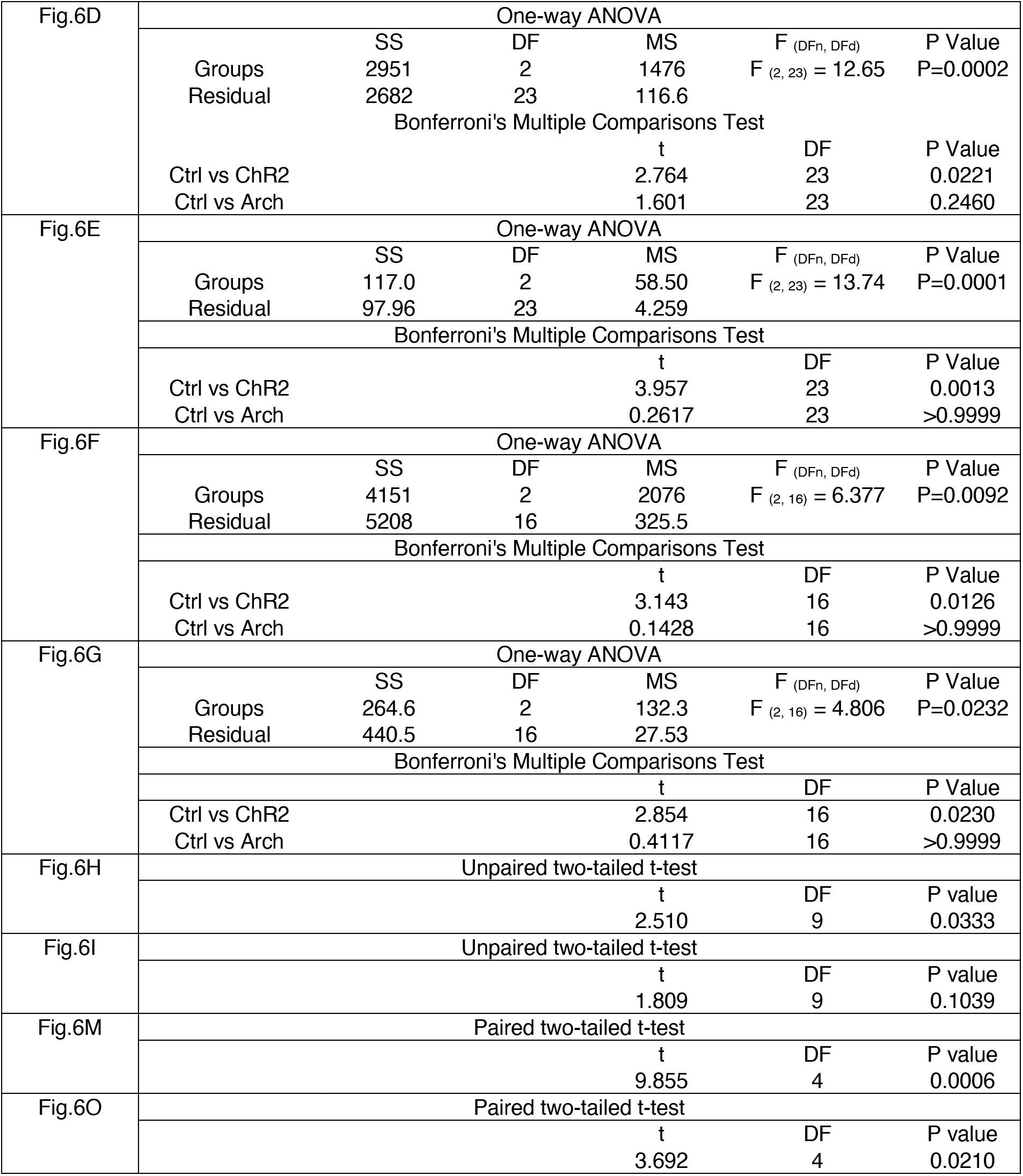
Statistical Information for Figure 6.

**Table S7.**
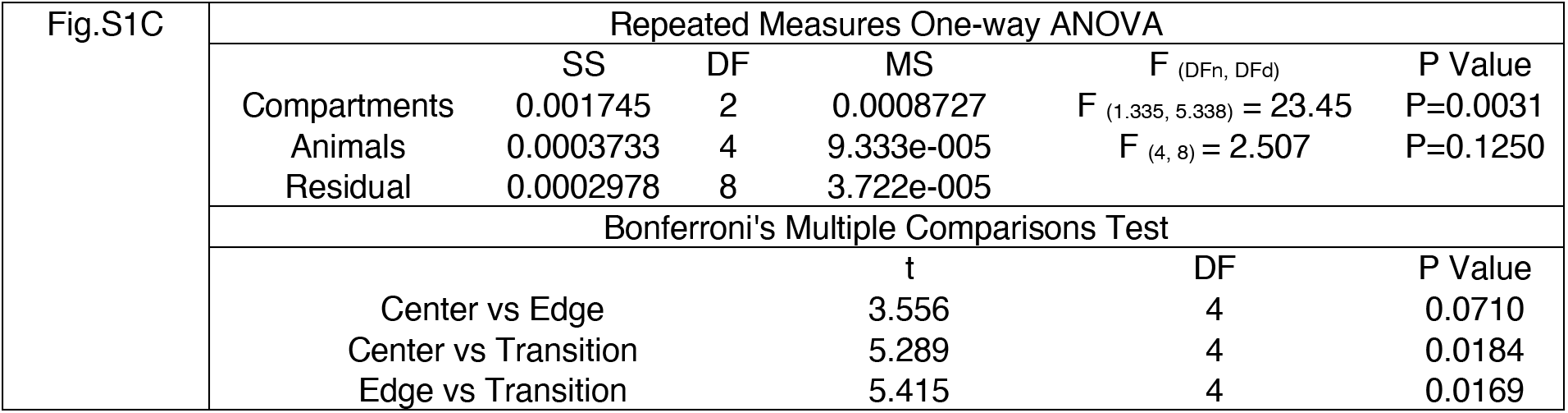
Statistical Information for Figure S1.

| Fig.S1C | Repeated Measures One-way ANOVA |  |  |  |  |  |
| --- | --- | --- | --- | --- | --- | --- |
|  |  | SS | DF | MS | F (DFn, DFd) | P Value |
|  | Compartments | 0.001745 | 2 | 0.0008727 | F (1.335, 5.338) = 23.45 | P=0.0031 |
|  | Animals | 0.0003733 | 4 | 9.333e-005 | F (4, 8) = 2.507 | P=0.1250 |
|  | Residual | 0.0002978 | 8 | 3.722e-005 |  |  |
|  | Bonferroni's Multiple Comparisons Test |  |  |  |  |  |
|  |  |  |  | t | DF | P Value |
|  | Center vs Edge |  |  | 3.556 | 4 | 0.0710 |
|  | Center vs Transition |  |  | 5.289 | 4 | 0.0184 |
|  | Edge vs Transition |  |  | 5.415 | 4 | 0.0169 |

**Table S8.** Statistical Information for Figure S3.

|  |  |  |  |
| --- | --- | --- | --- |
| Fig.S3N | Unpaired two-tailed t-test |  |  |
|  | t | DF | P value |
|  | 6.280 | 22 | <0.0001 |
| Fig.S3O | Unpaired two-tailed t-test |  |  |
|  | t | DF | P value |
|  | 4.538 | 22 | 0.0002 |
| Fig.S3P | Unpaired two-tailed t-test |  |  |
|  | t | DF | P value |
|  | 6.541 | 22 | <0.0001 |

**Table S9.**
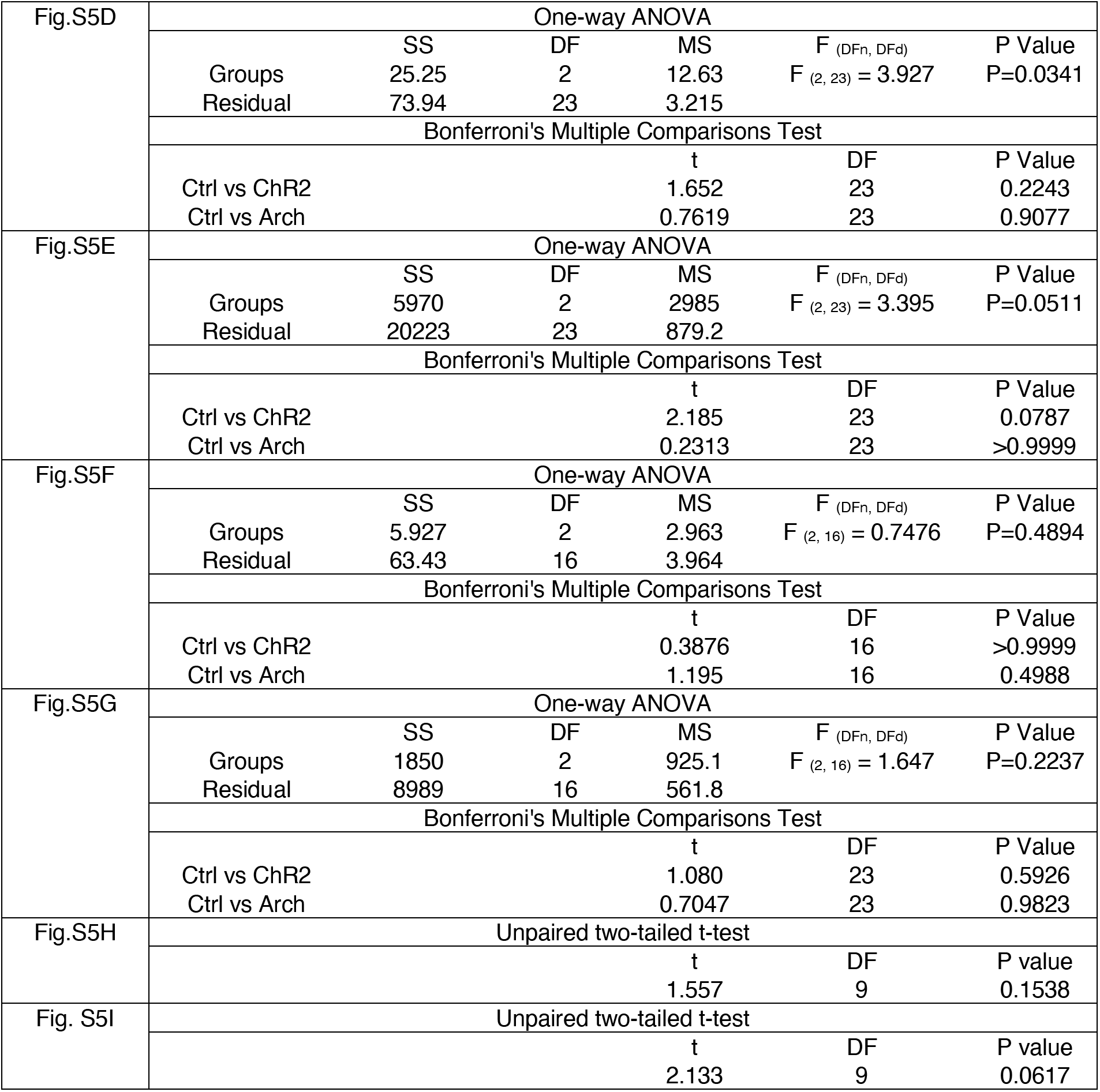
Statistical Information for Figure S5.

